# A ketogenic diet reduces age-induced chronic neuroinflammation in mice

**DOI:** 10.1101/2023.12.01.569598

**Authors:** Mitsunori Nomura, Natalia Faraj Murad, Sidharth S Madhavan, Brenda Eap, Thelma Y Garcia, Carlos Galicia Aguirre, Eric Verdin, Lisa Ellerby, David Furman, John C Newman

**Author notes:** (M.N.) (N.F.M.) (S.S.M.) (B.E.) (T.Y.G.) (C.G.A.) (E.V.) (L.E.) (D.F.).

## Abstract

Beta-hydroxybutyrate (BHB) is a ketone body synthesized during fasting or strenuous exercise. Our previous study demonstrated that a cyclic ketogenic diet (KD), which induces BHB levels similar to fasting every other week, reduces midlife mortality and improves memory in aging mice. BHB actively regulates gene expression and inflammatory activation through non-energetic signaling pathways. Neither of these activities has been well-characterized in the brain and they may represent mechanisms by which BHB affects brain function during aging. First, we analyzed hepatic gene expression in an aging KD-treated mouse cohort using bulk RNA-seq. In addition to the downregulation of TOR pathway activity, cyclic KD reduces inflammatory gene expression in the liver. We observed via flow cytometry that KD also modulates age-related systemic T cell functions. Next, we investigated whether BHB affects brain cells transcriptionally *in vitro*. Gene expression analysis in primary human brain cells (microglia, astrocytes, neurons) using RNA-seq shows that BHB causes a mild level of inflammation in all three cell types. However, BHB inhibits the more pronounced LPS-induced inflammatory gene activation in microglia. Furthermore, we confirmed that BHB similarly reduces LPS-induced inflammation in primary mouse microglia and bone marrow-derived macrophages (BMDMs). BHB is recognized as an inhibitor of histone deacetylase (HDAC), an inhibitor of NLRP3 inflammasome, and an agonist of the GPCR Hcar2. Nevertheless, in microglia, BHB’s anti-inflammatory effects are independent of these known mechanisms. Finally, we examined the brain gene expression of 12-month-old male mice fed with one-week and one-year cyclic KD. While a one-week KD increases inflammatory signaling, a one-year cyclic KD reduces neuroinflammation induced by aging. In summary, our findings demonstrate that BHB mitigates the microglial response to inflammatory stimuli, like LPS, possibly leading to decreased chronic inflammation in the brain after long-term KD treatment in aging mice.

## Introduction

The ketone body beta-hydroxybutyrate (BHB) is a small metabolite primarily synthesized in the liver that functions as a glucose-sparing energy source in extrahepatic organs, including the brain and heart^1–3^. In healthy adults, the non-fasting blood concentration ranges from 0.05-0.25 mM, while with prolonged exercise, fasting, or the absence of dietary carbohydrates, it can reach 0.5-8 mM^2^. In addition to its role as an energy source, BHB is increasingly recognized as a signaling metabolite. For instance, it acts as an inhibitor of NLRP3 inflammasome^4,5^, an inhibitor of class I histone deacetylases (HDACs)^6^, and also a modifier of protein lysine beta-hydroxybutyrylation (Kbhb)^7^. Moreover, BHB serves as a ligand for the G-protein-coupled receptor hydroxycarboxylic acid receptor 2 (Hcar2, HCA2, GPR109A)^8^. A recent study suggests that BHB can prevent vascular senescence by binding with hnRNP A1, a member of the hnRNP family responsible for RNA processing, and enhancing the expression of Oct4^9^. BHB, a chiral molecule, has two enantiomers: R-BHB and S-BHB. The former is the dominant endogenous product and can be readily catabolized into acetyl-CoA and ATP, while the latter cannot^2^.

A ketogenic diet (KD) contains high fat, moderate amounts of protein, and little to no carbohydrate to stimulate endogenous ketone body production, particularly BHB. We previously demonstrated that a non-obesogenic “cyclic” KD, alternated with a typical diet every other week, reduces mortality and enhances memory in aging mice^10^. During the study, the peak plasma levels of R-BHB in the cyclic KD-fed mice were 1.5-2.5 mM upon consuming the KD^10^. Independently, Roberts et al. showed that a non-obesogenic, isocaloric KD enhances longevity and health span, including brain function, in aging mice^11^. Recently, Tomita et al. demonstrated that feeding aging mice with the exogenous BHB precursor 1,3-butanediol increases lifespan^12^. Their findings suggest that promoting BHB production might directly enhance longevity and health in aging mice. The mechanisms by which KD and/or BHB enhance longevity are incompletely understood. One possibility is via regulation of the target of rapamycin (TOR) signaling, a pathway known to broadly regulate longevity, most profoundly by caloric restriction^13^. The network responds to nutrients (e.g., glucose, amino acids), and regulates cell metabolism, such as autophagy, protein synthesis, and lipid metabolism^14^. Rapamycin, a TOR inhibitor, extends the lifespan of various strains of mice^15^. In our study, we conducted RNA sequencing (RNA-seq) and RT-qPCR analyses and found that both a one-week KD and a one-year cyclic KD reduced TOR activation and upregulated PPARα target genes in the liver and kidney^10^. Roberts et al. also suggest an isocaloric KD inhibits hepatic TOR signaling^11^. While both studies indicate that the KD focuses on improving memory in the aging brain, the transcriptional changes in the brain after KD treatments and the cell-specific effects in the brain following BHB activation were not explored.

In this study, we first use RNA-seq from the liver of a one-year cyclic KD-fed cohort^10^, to show that a long-term non-obesogenic KD strongly inhibits hepatic chronic inflammation. Using *in vitro* models, we demonstrate that the acid form of R-and S-BHB significantly reduces the lipopolysaccharide (LPS)-induced inflammatory response in human and mouse primary microglia and bone marrow-derived macrophages (BMDMs). In contrast, R-BHB modestly activates inflammatory pathways in LPS-untreated primary human astrocytes and neurons. Examining the brain gene expression of our KD cohorts^10^, we found that one-year KD feeding reduces age-induced neuroinflammation, a distinct pattern from the one-week KD feeding. In summary, BHB decreases LPS-induced inflammation in microglia and reduces brain inflammation in aging *in vivo*, supporting a key role for inflammatory modulation in the effect of KD on ameliorating brain aging phenotypes in mice.

## Materials and Methods

### Mouse strains, housing, and husbandry

The mouse experiments involved mice housed either at Buck Institute for primary cell isolation or previously housed as described at Gladstone Institutes for long-term cyclic KD studies^10^. All mice were maintained according to the National Institutes of Health guidelines, and all experimental protocols were approved by the Buck Institute Institutional Animal Care and Use Committee (IACUC) or the UCSF IACUC (for mice previously housed at Gladstone). All mice were maintained in a specific pathogen-free barrier facility on a 7:00 am to 7:00 pm light cycle (6:00 am to 6:00 pm at Gladstone). To prepare primary microglia, astrocytes, and bone marrow-derived macrophages (BMDMs), we initially acquired C57BL/6 mice from the National Institute on Aging’s Aged Rodent Colony. The mice were kept in an animal facility and maintained under a light cycle from 6:00 a.m. to 6:00 p.m. We ensured that all mice were managed following the National Institutes of Health guidelines, and every procedure and protocol was approved by the Buck Institutional Animal Care and Use Committee (IACUC). The details of all mice using the *in vivo* RNAseq experiments and immunophenotyping in blood are described previously^10^. Briefly, C57BL/6 male mice were obtained from the National Institute on Aging’s Aged Rodent Colony at 11 months of age and started on experimental diets at 12 months of age. Aging mice were monitored with increasing frequency based on a protocol developed in collaboration with UCSF LARC veterinarians. The groups of mice designated for tissue collection after 1 week and 12 months on the experimental diets were separate from the other lifespan and behavioral testing cohorts.

### Mouse diets and feeding

The mouse diet specifics have been previously described^10^. Upon arrival, until switched to experimental diets, mice were fed the Gladstone Institutes standard vivarium chow (chow) with 24 % protein, 13 % fat, and 62 % carbohydrates. The customized diets from Envigo contained the following macronutrient content per calorie: Control (CD) with 10% protein, 13% fat, and 77% carbohydrate (TD.150345); KD with 10% protein and 90% fat (TD.160153). Both diets had similar micronutrient content, fiber, and preservatives on a per-calorie basis, had similar fat sources, and were always provided *ad libitum*. All food was changed once per week; for the cyclic KD condition, this involved switching diets.

### RNA-Seq

The dataset of one-week KD liver RNA-seq has been already published^10^. For in vivo sequencing, RNA was isolated using the Direct-zol RNA MiniPrep kit (Zymo Research) following the manufacturer’s protocol. Subsequent RNA sample processing was conducted by UC Davis DNA Technologies & Expression Analysis Core. RNA integrity and concentrations were assessed using an Agilent Bioanalyzer. The library was prepared using the KAPA RNA HyperPrep Kit (Roche) and QIAseq FastSelect RNA Removal Kit (QIAGEN), and sequencing was performed on a NovaSeq 6000 (Illumina). For in vitro (3’ Tag-Seq) sequencing, RNA was isolated using the Quick-RNA MicroPrep Kit (Zymo Research) according to the manufacturer’s protocol. Downstream processing of the RNA samples was carried out by the UC Davis DNA Technologies & Expression Analysis Core. RNA integrity and concentrations were confirmed using an Agilent Bioanalyzer. The library was prepared with the KAPA RNA HyperPrep Kit (Roche) and sequenced on a HiSeq 4000 (Illumina). Quality reports were assessed before and after trimming reads using FastQC (http://www.bioinformatics.babraham.ac.uk/projects/fastqc/) and MultiQC (https://multiqc.info). For the analysis, low-quality bases and adapters were removed using trimmomatic and reads were mapped to the reference genome (GRCm38, http://nov2020.archive.en-sembl.org/Mus_musculus/Info/Index) with STAR^16,17^. The count matrixes were generated with RSEM^18^. For differential expression analysis, DEseq2 was used^19^, and p-values were corrected using False Discovery Rate (FDR). For the differentially expressed genes (DEGs), we considered adjusted p-value < 0.05 and |log fold change| > 1.0. Panther database was used for transcripts annotation^20^. Enrichment analyses to find significant pathways/signatures associated with the DEGs were done using clusterprofiler^21,22^, DOSE^23^, org.Hs.eg.db^24^, and org.Mm.eg.db^25^. Data were visualized using packages ggplot2^26^ and gridExtra. Gene lists were prepared for gene set enrichment analysis (GSEA)^27^ by ranking by t-statistic and duplicate or missing gene identifiers were removed. We used the Molecular Signatures Database (MSigDB) hallmark gene set, gene ontology (GO)^28^, and Kyoto Encyclopedia of Genes and Genome (KEGG)^29^ analyses. GSEA for the gene ontology (GO) biological process was executed for genes with a p-value cutoff of 0.01 and GSEA for the KEGG pathway was executed for genes with a p-value cutoff of 0.05. To visualize concordant and discordant gene overlap, the Rank-Rank Hypergeometric Overlap (RRHO) package was used^30^. We selected the lists of common genes between the pairs of comparisons and listed by down-and up-regulated. We ordered all the genes by log-foldchange and generated the gene lists that provided the most significant overlap.

### Flow cytometry

For the blood samples, red blood cells were lysed using an ammonium-chloride-potassium (ACK) buffer. Lymphocytes from the blood were stained with the LIVE/DEAD Fixable Green Dead Cell Stain Kit (L34970; Invitrogen) and antibodies against the surface antigens: TCRβ (H57-597, BioLegend), CD4 (RM4-5, BioLegend), CD8 (53-6.7, BioLegend), PD-1 (J43; BD Biosciences), and then fixed with 1% PFA. For the Foxp3 staining, the cells underwent an additional staining process using anti-Foxp3 (MF23; BD Biosciences) following the manufacturer’s protocol. Stained cells were acquired on an LSRII (BD), and the FlowJo program was used for data analysis.

### Cell culture

Primary human microglia (#1900), astrocytes (#1800), and neurons (#1520) were purchased from ScienCell. No identification information such as sex, age, or origin of the cells was provided. The culture media were Microglia Medium (#1901), Astrocytes Medium (#1801), and Neuron Medium (#1521), respectively (all include 10 % FBS, penicillin-streptomycin, and growth supplement). For the preparation of primary mouse microglia and astrocytes, postnatal 0-2 pups (C57BL/6 background) were euthanized by decapitation. The brains were collected and triturated in PBS using a serological pipette. After centrifugation, cells were resuspended and plated in poly-L-lysine-coated flasks. The cells were cultured in DMEM (10-013-CV; Corning) with 10 % FBS (35-011-CV; Corning) and penicillin-streptomycin (30-002-CI; Corning) and the media were replaced at two to three days a week. After ten days, microglia were removed and collected by tapping and shaking the flasks. The remaining cells in the flasks were used as astrocytes. Both IMG and BV-2 cells were kindly provided by the Andersen Lab at the Buck Institute and cultured in DMEM with 10 % FBS and penicillin-streptomycin. For the preparation of bone marrow-derived macrophages, adult C57BL/6 mice were euthanized, and the leg bones were cleaned. The bone marrow was flushed from the bones using PBS. The cells were then centrifuged, resuspended, and plated on Petri dishes. The cells were cultured in DMEM with 10% FBS, penicillin-streptomycin, and 10ng/ml of M-CSF (576404; BioLegend). New media were added three days later, and after seven days, macrophages were collected for subculturing. Cells were treated with 10 mM (several concentrations were used in Supplemental Figure 7B) R-BHB (54920; Sigma), 10 mM S-BHB (54925; Sigma), 10 mM Na-R-BHB (298360; Sigma), 10 mM Na-S-BHB (sc-236887; Santa Cruz), 5 mM Bu (L13189; Alfa Aesar), 5 mM Na-Bu (B5887; Sigma), 100 ng/ml LPS (sc-3535; Santa Cruz), 10 mM NaCl (BDH9286; VWR), 5 mM ATP (tlrl-atpl; InvivoGen), 10 μM nigericin (tlrl-nig; InvivoGen), 1 μM MCC950 (17510; Cayman) following the described time course.

### Small interfering RNA (siRNA) transfection

Three synthetic siRNAs, siGENOME Non-Targeting siRNA Control Pool #1 (siCtrl; D-001206-14-05), siGENOME Mouse Hcar2 siRNA (siHcar2; M-040890-00-0005), and siGENOME Mouse Hnrnpa1 siRNA (siHnrnpa1; M-040887-01-0005), were purchased from Horizon Discovery. The oligos were transfected into primary mouse microglia using INTERFERin (101000028; Polyplus) following the manufacturer’s protocol. The cells were used for further experiments after 24 hours.

### Quantitative PCR

RNA was isolated by Quick-RNA MicroPrep Kit (R1051; Zymo Research) and cDNA synthesis was carried out with Superscript cDNA Synthesis Kit (1708891; BioRad) and real-time quantitative PCR was performed using iTaq Universal SYBR Green Supermix (1725121; BioRad) in a BioRad CFX96 Real-Time System. Gene expression analyses were normalized to the B2m housekeeping gene. Primers; B2m: Fw_ACAGTTCCACCCGCCTCACATT, Rv_TAGAAAGACCAGTCCTT- GCTGAAG; Il1b: Fw_TGGACCTTCCAGGATGAGGACA, Rv_GTTCATCTCGGAGCCTG- TAGTG; Il6: Fw_TACCACTTCACAAGTCGGAGGC, Rv_CTGCAAGTGCATCATCGTTGTTC; Ccl2: Fw_GCTACAAGAGGATCACCAGCAG, Rv_GTCTGGACCCATTCCTTCTTGG; Hcar2: Fw_ CTGTTTCCACCTCAAGTCCTGG, Rv_ CATAGTTGTCCGTCAGGAACGG; Hnrnpa1: Fw_ CGAAACAACCGACGAGAGTCTG, Rv_ CATGGCAGCATCCACTTCTTCC.

### Western blotting

Cells and tissues were lysed by a RIPA buffer (1% NP-40, 1% sodium deoxycholate, 0.1% SDS,150mM NaCl, and 25mM Tris-HCl, pH 7.6) with protease inhibitor cocktail (11836170001; Roche). The proteins of the supernatant for the IL-1β secretion inflammasome assay were precipitated by chloroform and methanol. These lysates were prepared by a sample buffer (NP0007; Invitrogen) with 10 % 2-Mercaptoethanol and loaded onto 4-20% precast polyacrylamide gels (4561096, 5671095; Bio-Rad). Imaging was performed with enhanced chemiluminescent detection (34096; Thermo Scientific) on an Azure Biosystems c600 imager. Antibodies; anti-H3K9bhb: PTM-1250, PTM Biolabs; anti-H3K9ac: #9496, Cell Signaling; anti-H3: 07-690, Millipore; anti-Gapdh: 60004-1-lg, Proteintech; anti-HSP90: 610419, BD Biosciences; anti-Kac: #9441, Cell Signaling; anti-Kbhb: PTM-1201, PTM-Biolabs; anti-IL-1β: GTX74034, GeneTex; anti-mouse IgG: #7076, Cell Signaling; anti-rabbit IgG: #7074, Cell Signaling.

### IL-1β ELISA

IL-1β secretion was detected by IL-1 beta Mouse Uncoated ELISA Kit (88-7013-88; Invitrogen) following the manufacturer’s protocol.

### Quantification and statistical analysis

Figures are presented as mean ± SD, and P values are calculated by one-way ANOVA with additional post-hoc testing for group comparisons. In the figures, P values are denoted by real values or asterisks (*P < 0.05, **P < 0.01, ***P < 0.001, ****P < 0.0001). Statistical calculations are carried out in Prism 10 (GraphPad). In Supplemental Figure 2B, one value was identified as an outlier by both the ROUT and Grubb’s test and removed.

## Results

### One-year KD reduces hepatic chronic inflammation

Previously, we demonstrated that an alternate-week cyclic KD, which is non-obesogenic, can enhance the lifespan and healthspan in aging mice^10^. We and Roberts et al. also showed that non-obesogenic KDs suppress TOR activity in the liver^10^. Goldberg et al. found that a one-week KD activates γδ T cells in adipose tissue and restricts inflammation, whereas a four-month ad libitum KD promotes obesity and adipose chronic inflammation in mice^31^, which suggests the effects of a KD in chronic inflammation depend on the feeding regimens (e.g., durations, obesogenic / non-obesogenic). Chronic inflammation is a hallmark of aging and is associated with morbidity and mortality in aging^14,32,33^. We hypothesized a KD inhibits chronic inflammation during aging (“inflammaging”) if the KD does not cause obesity. To examine changes in the liver transcription after a long-term (one-year) cyclic KD treatment, we conducted bulk RNA-seq and Gene Set Enrichment Analysis (GSEA) (Figure 1A). The tissue samples were collected during a control diet (CD) feeding period to focus on persistent long-term effects related to aging rather than transient acute effects of the KD. We found many KD-regulated genes (Supplemental Figure 1A), and the KD led to the suppression of “mTORC1 signaling” and “PI3K AKT mTOR signaling”, as expected (Figure 1B). The cyclic KD also led to the suppression of several hallmark pathways, including “TNFA Signaling via NFkB,” “Interferon Alpha Response,” “Inflammatory Response,” and “Interferon Gamma Response”, which are generally related to chronic inflammation (Figure 1B). Gene Ontology (GO) and Kyoto Encyclopedia of Genes and Genomes (KEGG) analyses revealed that the KD suppresses immune response-related pathways such as the “immune system process” (GO), and “Phagosome” (KEGG) (Supplemental Figure 1B, C). However, a one-week KD did not change the inflammatory pathways (via reanalysis of a data set reported previously^10^) (Supplemental Figure 1D). These findings suggest that, contrary to short-term feeding, a non-obesogenic KD can mitigate chronic inflammation in the liver over an extended period.

**Figure 1.**
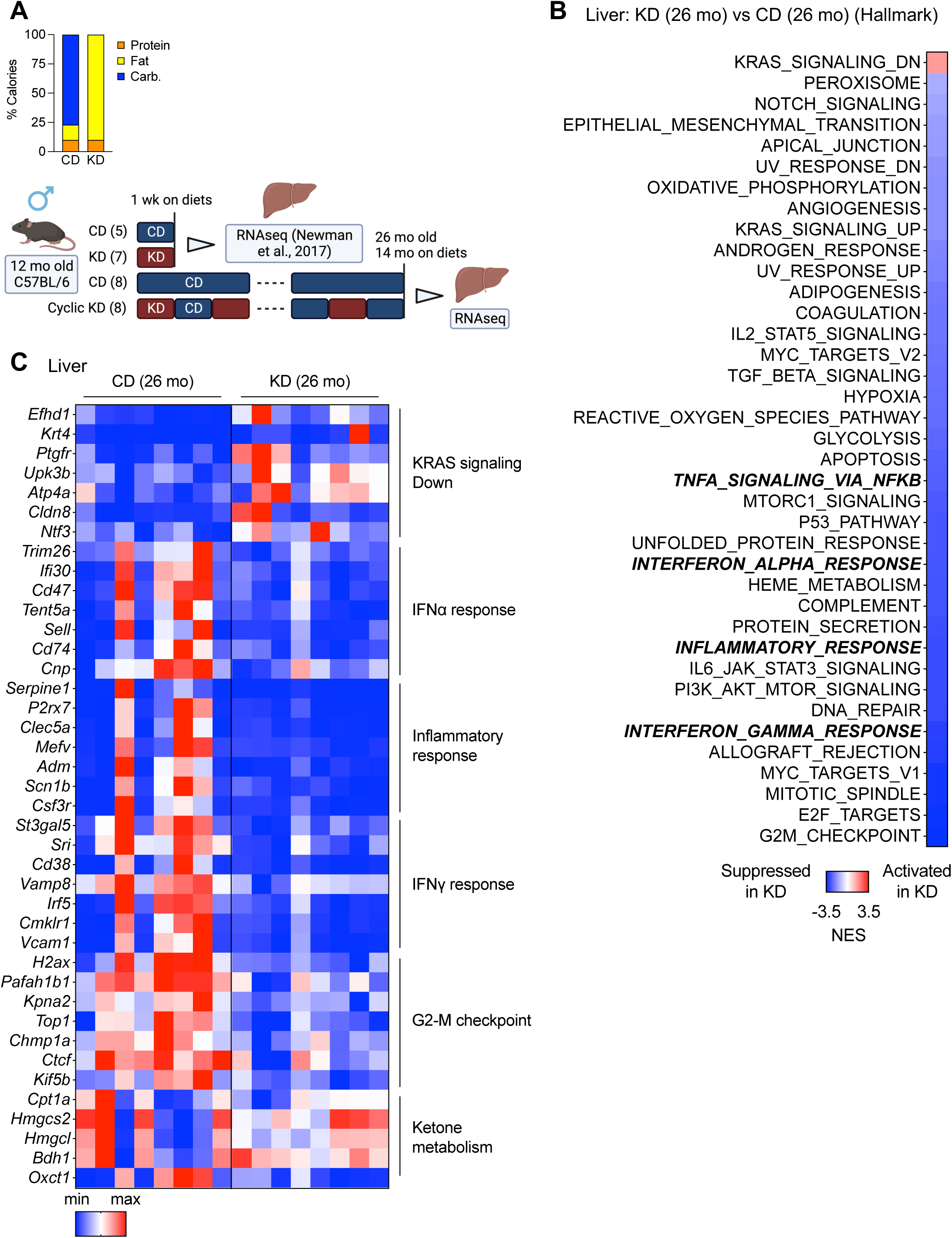
One-year KD reduces hepatic chronic inflammation. (A) Diet composition and experimental timeline with mouse numbers. An analysis of RNA-seq after the one-week KD was published before^10^. (B) GSEA (hallmark) analysis (KD vs. CD) of 26-month liver on diets for 14 months, collected at a dark cycle during CD-fed week (n=8 mice per group). (C) Heatmap of gene expressions in the liver.

Recent studies suggest that KDs and BHB can improve T cell immune functions in both humans and mice^34–36^. To examine the impact of a cyclic KD on age-related immunophenotyping systemically, blood samples were collected and analyzed after a one-year KD intervention at 24 months of age (Supplemental Figure 2A). As with the liver, these samples were collected during a period of feeding on a CD. We compared aging effects by using 3-month and 12-month-old mice. The KD did not significantly alter the overall proportion of either CD4^+^ or CD8^+^ cells; however, it significantly reversed the CD8^+^/CD4^+^ ratio that was increased by the aging process (Supplemental Figure 2B). The diet did not affect the populations of Foxp3^+^/CD4^+^ (regulatory T cells), PD-1^+^/CD4^+^, and PD-1^+^/CD8^+^ (Supplemental Figure 2C, D). These findings indicate that a non-obesogenic KD could partially reverse systemic immune functions affected by aging.

**Figure 2.**
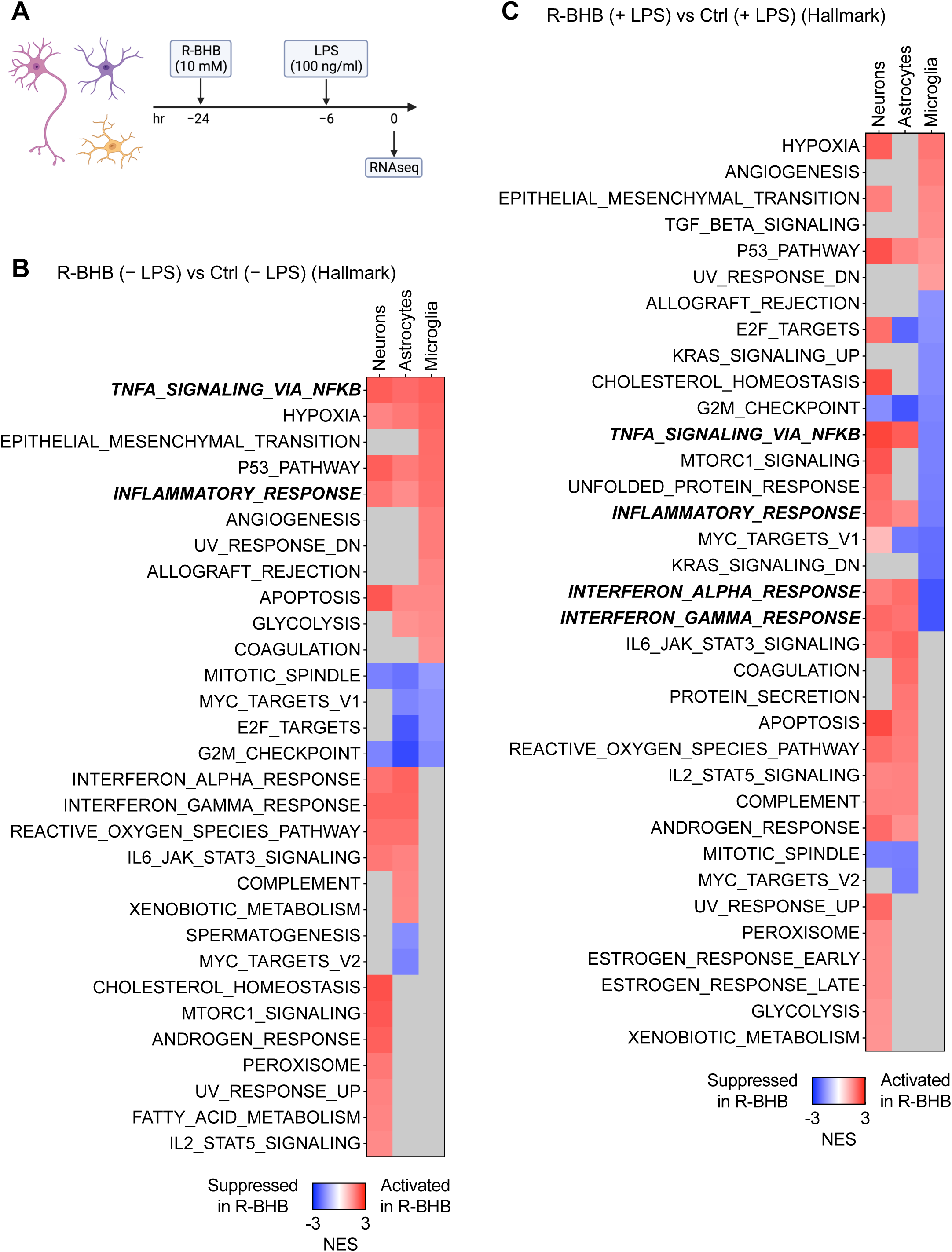
R-BHB reduces LPS-induced inflammation in human primary microglia. (A) Experimental timeline. (B, C) GSEA (hallmark) analysis (R-BHB vs. Ctrl) without (B) or with (B) LPS treatment in human primary neurons/astrocytes/microglia (n=3 per group). Gray squares: not significant.

### R-BHB reduces LPS-induced inflammation in human primary microglia

In mice, KD induces hepatic R-BHB production and increases plasma and brain levels by more than tenfold^37^. We hypothesize that R-BHB directly modulates brain functions, including chronic neuroinflammation. To examine the transcriptional effects of R-BHB on brain cells, we initially employed *in vitro* culture models for primary human microglia, astrocytes, and neurons. Chronic inflammation is also a hallmark of brain aging and neurodegenerative diseases^38,39^. There are no established *in vitro* models that precisely represent age-induced inflammation, but to simulate systemic inflammation *in vitro*, we used LPS, which causes brain inflammation in *in vivo* models^40^. Our preliminary experiment demonstrated that a six-hour LPS incubation most strongly induces inflammatory cytokine levels in the microglia (Supplemental Figure 3A). The total incubation time for 10 mM R-BHB acid, the endogenous form of BHB induced by ketosis, was 24 hours, with activation by LPS for the final six hours. Gene expression was analyzed by RNA-seq (Figure 2A). Volcano plots indicate that R-BHB substantially altered gene expression in all three cell types, regardless of LPS stimulation. Meanwhile, LPS had the strongest effect on gene expression in microglia, but only demonstrated a weak effect on astrocytes and little effect on neurons (Supplemental Figure 3B-D). Hallmark analysis and gene expression patterns indicate that, in the absence of LPS stimulation, R-BHB upregulates inflammatory signaling through “TNFA Signaling via NFkB” and “Inflammatory Response,” while inhibiting cell cycle regulation pathways via “Mitotic Spindle” and “G2-M Checkpoint” in all three types of cells (Figure 2B and Supplemental Figure 4A-C). LPS stimulation activates inflammatory signaling in microglia and astrocytes, but not in neurons (Supplemental Figure 3E). With LPS co-treatment, R-BHB still activates inflammation pathways in neurons and astrocytes, but it suppresses signaling that includes the “Interferon Alpha Response” and “Interferon Gamma Response” in microglia (Figure 2C, Supplemental Figure 4B-D). Indeed, BHB substantially blunted the overall inflammatory gene response of LPS in microglia (Figure 2C and Supplemental Figure 4C). LPS stimulation reduced the expression of genes involved in ketogenesis, specifically CPT1A, HMGCL, and BDH1 in microglia and R-BHB induced certain monocarboxylate transporters (MCTs) in a cell-dependent manner. For instance, SLC16A7 expression was increased in neurons, and SLC16A6/10/14 was increased in microglia (Supplemental Figure 4E), which suggests a cell-specific ketogenic capacity in the brain. Based on GO and KEGG analyses, R-BHB appears to suppress several key pathways, including the “innate immune response” (GO) and “Chemokine signaling pathway” (KEGG) following LPS stimulation in microglia but not neurons and astrocytes (Supplemental Figures 5A-C and 6A-C). In summary, R-BHB alters the transcriptome in a brain cell-dependent manner, with specific antiinflammatory effects on microglia.

### BHB acids reduce LPS-induced inflammation in mouse primary microglia

Next, we investigated the dose-dependent anti-inflammatory effects of R-BHB in primary mouse microglia, following the same time course as described in Figure 2A (Figure 3A). R-BHB (5 and 10 mM) effectively suppressed LPS-induced inflammatory cytokines, including IL-1β (Il1b), IL-6 (Il6), and TNFα (Tnf), in the cells, whereas the chemokine (C-C motif) ligand 2 (Ccl2) was unaffected. No effect was observed with a concentration of 1 mM (Supplemental Figure 7B). There are various possible ways in which R-BHB may regulate inflammation in microglia, such as inhibiting energy supply^41^, HDAC inhibition^42^, regulation of Kbhb^7^, the NLRP3 inflammasome^4,43^, Hcar2 (also known as HCA2 and GPR109A) activation^44,45^, and interaction with hnRNA A1^9^.

**Figure 3.**
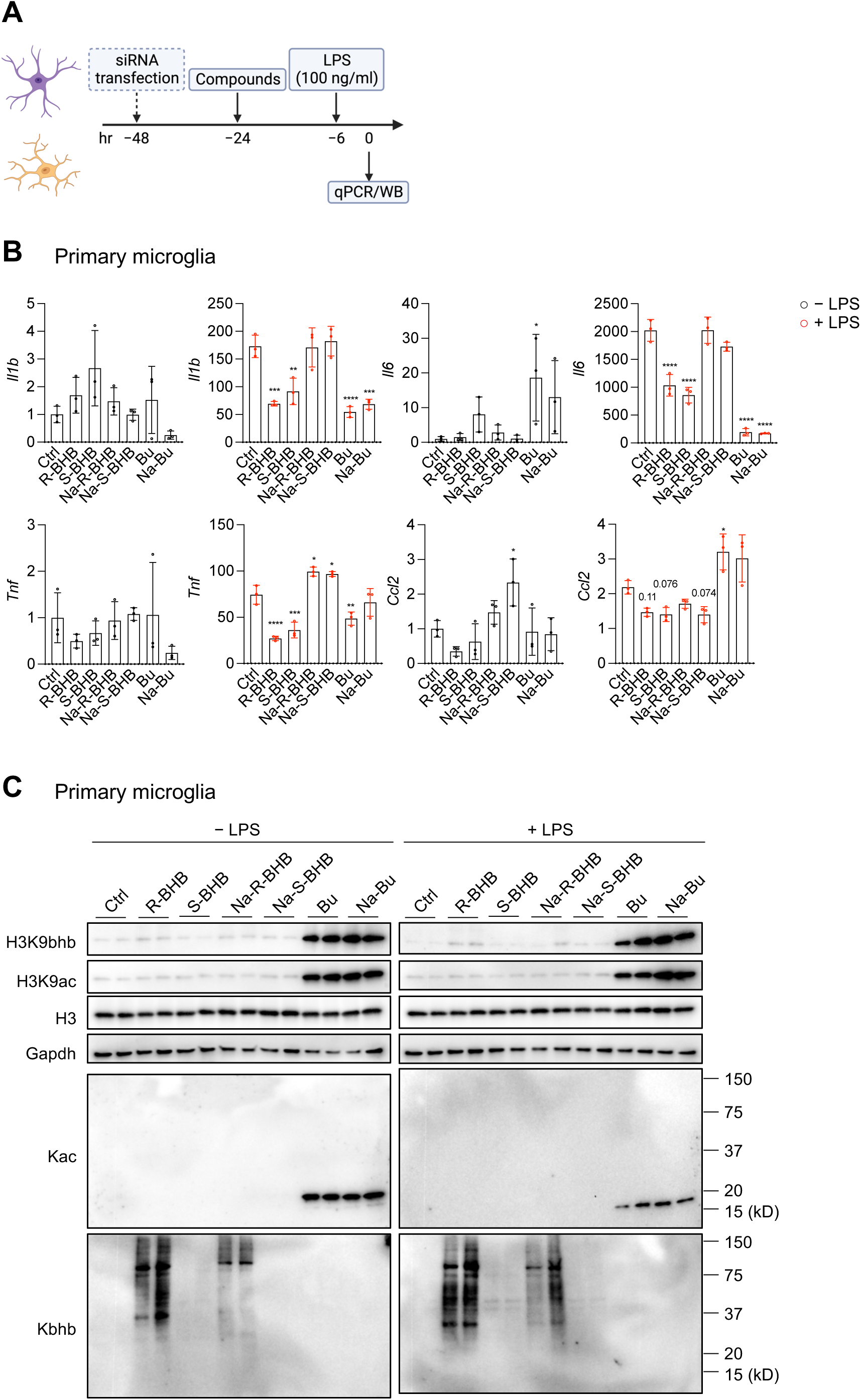
BHB acids reduce LPS-induced inflammation in mouse primary microglia. (A) Experimental timeline. (B) mRNA expression in mouse primary microglia (n=3 per group). All data are presented as mean ± SD. One-way ANOVA with Dunnet’s correction for multiple comparisons. Compare the mean of each sample with the mean of Ctrl. (C) Protein expression in mouse primary microglia (n=2 per group). All data are representative of two independent experiments. Abbreviations: R-BHB, R-BHB acid; S-BHB, S-BHB acid; Na-R-BHB, R-BHB salt; Na-S-BHB, S-BHB salt; Bu, butyric acid; Na-Bu, butyrate (salt).

To examine the mechanisms, we used acid and sodium forms of both BHB enantiomers (R-BHB and S-BHB), each at a concentration of 10 mM, along with the related structures to BHB, such as sodium butyrate and butyric acid, which are also HDAC inhibitors^46^, at a concentration of 5 mM. We administered acid forms (R-BHB, S-BHB, and butyric acid (Bu)) and sodium salt forms (Na-R-BHB, Na-S-BHB, sodium butyrate (Na-Bu)) (Supplementary Figure 7A). Upon adding these compounds to the culture media, all acid forms caused a small decrease in the media pH, while the salt forms did not (Supplementary Figure 7A). R-BHB, S-BHB, Bu, and Na-Bu, but not Na-R-BHB and Na-S-BHB, decreased the expression of inflammatory cytokines (Il1b, Il6, and Tnf) when treated with LPS (Figure 3B). In primary microglia, the inclusion of sodium chloride (10 mM NaCl) does not alter the anti-inflammatory responses by R-BHB acid (Supplementary Figure 7C), suggesting sodium salt by itself does not affect inflammatory regulation. These findings indicate the mechanism(s) by which BHB reduces inflammation may be highly structurally specific or be modulated by small changes in the local cellular environment.

Since R-BHB serves as an endogenous HDAC inhibitor^6^ and a regulator of Kbhb^7^, we identified protein lysine acetylation (Kac) and Kbhb by western blotting (Figure 3C). Subsequently, we analyzed how these compounds induce epigenetic modifications. Without LPS, R-BHB and Na-R-BHB significantly stimulated the expression of protein Kbhb (the antibody is R form specific). Additionally, both compounds showed a mild increase in histone 3 lysine 9 beta-hydroxybutyration (H3K9bhb) while not influencing Kac or histone 3 lysine 9 acetylation (H3K9ac). On the other hand, both Bu and Na-Bu substantially increased the levels of H3K9bhb, H3K9ac, and Kac but not Kbhb. The same trend was observed in the presence of LPS as well (Figure 3C). The findings imply that BHB does not have a direct effect on HDAC inhibition to impede inflammation in microglia. BHB activates the GPCR Hcar2, which is reported to be necessary for BHB’s neuroprotective effect^44,45^. BHB also directly binds to heterogeneous nuclear ribonucleoprotein A1 (hnRNA A1) and upregulates Oct4, a regulator of pluripotency, in both vascular smooth muscle and endothelial cells^9^. We utilized small interfering RNA (siRNA) targeting Hcar2 and Hnrnpa1 to examine the respective functions of these genes in primary microglia (Supplemental Figure 7D), Both siRNAs decreased each expression by approximately 50% and LPS incubation led to a 100-fold rise in Hcar2 expression. Knockdown of either Hcar2 or Hnrnpa1 did not affect the anti-inflammatory effects of R-BHB. (Supplementary Figure 7D). These findings suggest that BHB’s anti-inflammatory effects are not directly attributable to Hcar2 activation or Hnrnpa1 binding in (at least) mouse primary microglia. Furthermore, we investigated whether BHBs inhibit the NLRP3 inflammasome in primary microglia (Supplemental Figure 8A). In the experiments, we used MCC950, an NLRP3 inflammasome inhibitor, as a positive control. All BHB compounds had no effect on IL-1β secretion after ATP-induced NLRP3 inflammasome activation (Supplemental Figure 8B, C). Surprisingly, the acid forms of R-and S-BHB, along with Bu and Na-Bu, increased nigericin-induced NLRP3 inflammasome activation (Supplemental Figure 8B, C). The modulation of the NLRP3 inflammasome by BHB probably depends on the cell types and activators present. These results suggest the mechanisms of how R-BHB acid inhibits inflammation in primary microglia are still elusive.

Next, we investigated the cell-specific effects of different forms of BHBs in mouse primary astrocytes (Supplemental Figure 9A, B), two microglia cell lines (IMG^47^ (Supplemental Figure 10A-C) and BV-2^48^ cells (Supplemental Figure 11A-C)), and bone marrow-derived macrophages (BMDMs) (Supplemental Figure 12A-C). In primary astrocytes, R-BHB and S-BHB demonstrated a reduction solely in Tnf expression (Supplemental Figure 9A), and their Kac and Kbhb expression patterns were equivalent to those observed in primary microglia (Supplemental Figure 9B). In IMG cells, the acid forms of BHB were observed to increase inflammation (Supplemental Figure 10A) and activate the NLRP3 inflammasome (Supplemental Figure 10B). BHBs increase the levels of both H3K9bhb and H3K9ac (Supplemental Figure 10C). In BV-2 cells, BHBs did not alter inflammatory expressions and did not activate NLRP3 inflammasome induced by ATP and nigericin (Supplemental Figure 11A, B). The acid forms of BHB induced H3K9bhb activation, but not H3K9ac (Supplemental Figure 11C). In BMDMs, the acid forms of BHB reduced inflammation (Supplemental Figure 12A). These forms induced nigericin-induced NLRP3 inflammasome (Supplemental Figure 12B). In the cells, both acid and salt forms of R-BHB induced H3K9bhb levels but not H3K9ac (Supplemental Figure 12C). In summary, these results indicate that BHB acids consistently decrease LPS-induced inflammation in primary microglia but that the effects on inflammation vary in other cell types.

### One-year KD reduces neuroinflammation distinct from one-week KD

Because a KD induces brain BHB levels more than 10-fold^37^, we hypothesize KDs may mitigate neuroinflammation in mouse *in vivo* models possibly through microglial functions. To explore the impact of a one-week and a one-year cyclic KD on transcriptional expression in the brain, we conducted an RNA-seq analysis on whole brain samples from our published cohorts^10^ (Figure 4A). As Figure 1A, the tissue samples were collected during a CD feeding. We found that the one-week KD, aging process, and one-year KD each altered the expression of numerous genes (Supplemental Figure 13A). The one-week KD induces several hallmark pathways such as “Hedgehog Signaling”, “Notch Signaling”, and “Inflammatory Signaling” in the brain (Supplementary Figure 13B). We observed an induction of genes related to ketone metabolism during the one-week KD (Supplemental Figure 13E), yet Kac and Kbhb levels were not detected in the brain after the one-week KD (Supplemental Figure 13C). The brains of 26-month-old mice fed a CD exhibited age-induced inflammation compared to 12-month-old mice on a CD (Supplemental Figure 13D, E). The one-year cyclic KD decreases inflammation-related pathways induced by aging, such as “TNFA Signaling via NFkB,” “Inflammatory Response,” and “Interferon Gamma Response,” in the brain (Figure 4B, C, and Supplemental Figure 13E). A GO analysis shows that the one-week KD activates angiogenesis-related functions (“angiogenesis”, “blood vessel morphogenesis”), which are inhibited by the one-year cyclic KD (Supplemental Figure 14A-C). A KEGG analysis shows that the one-year KD suppresses “Coronavirus disease – COVID-19”, which is activated by aging (Supplemental Figure 14D-F). These results demonstrate that the one-year KD lowers age-induced neuroinflammation, which differs from the impact of an acute (one-week) KD treatment.

**Figure 4.**
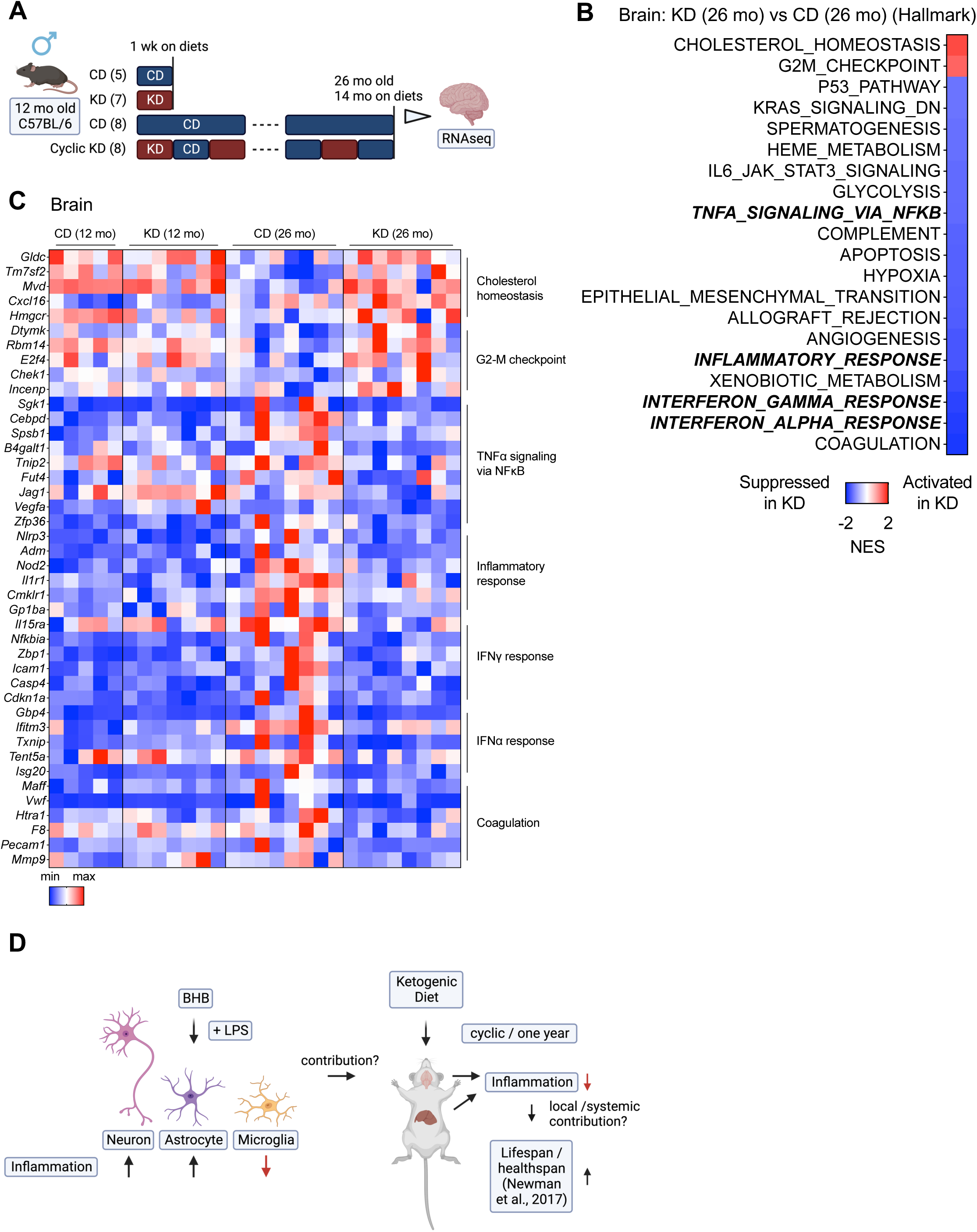
One-year KD reduces neuroinflammation distinct from one-week KD. (A) Experimental timeline with mouse numbers. (B) GSEA (hallmark) analysis (KD vs. CD) of 26-month brain on diets for 14 months, collected at night during CD-fed week (n=8 mice per group). (B) Heatmap of gene expressions in the brain. (D) Graphical abstract of the study.

We employed a Rank-Rank Hypergeometric Overlap (RRHO) analysis to identify overlapping genes that were altered in the same or opposite direction in two datasets^30^. In the brain, aging and the cyclic KD exhibit gene expression patterns that are almost entirely discordant (Supplemental Figure 15A), while the expression patterns of cyclic KD across tissues in the brain and liver are highly concordant, although they do not overlap completely (Supplemental Figure 15B). We next carried out a comparison between *in vitro* cell datasets (Figure 2) and *in vivo* brain datasets (Figure 4). In the absence of LPS, R-BHB (*in vitro*) and the one-week KD (*in vivo*) demonstrate discordant patterns in astrocytes and neurons, but concordant patterns in microglia (Supplemental Figure 15C). Recent *in vivo* analysis in mouse brains suggests that many of the genes implicated in the immune response, which were upregulated with age, were also upregulated by LPS^40^. While there is little concordance/discordance between LPS (*in vitro*) and aging (*in vivo*) patterns across all cells (Supplemental Figure 15D), R-BHB with a LPS treatment *in vitro* and cyclic KD *in vivo* yield concordant gene expression signatures in microglia but exhibit discordant expression in astrocytes and neurons (Supplemental Figure 15E). These findings suggest that microglial gene expression patterns detected *in vitro* align more closely with *in vivo* brain expression signatures than those of astrocytes and neurons.

## Discussion

Here we show that a cyclic KD inhibits chronic inflammation in mouse liver (Figure 1). R-BHB reduces inflammatory signaling in LPS-activated human primary microglia (Figure 2). We also confirmed that the acid forms of R-and S-BHB inhibit the inflammatory response in mouse primary microglia, by a mechanism yet to be identified (Figure 3). Finally, a cyclic KD reduced age-induced chronic inflammation in the mouse brain (Figure 4). A summary of this study is shown in Figure 4D.

In our RNA-seq experiments on human primary brain cells (Figure 2), we used 10 mM R-BHB acid. The acid form of R-BHB is a natural endogenous form of BHB^2^. Our research showed that the average plasma R-BHB levels for mice (3-, 12-, 22-months old, both sexes) fed either a one-week CD or a KD are roughly 0.2-0.4 mM and 2-4 mM, respectively. The average concentration of R-BHB in the brain is roughly 75 pmol/mg tissue (CD-fed) and 1,250 pmol/mg tissue (KD-fed)^37^. This suggests that during ketosis, such as with a KD diet, the brain may be exposed to higher concentrations of R-BHB compared to plasma R-BHB levels (15-20 times higher, compared to a standard diet), and a physiological range of 10 mM R-BHB is present in the brain under ketosis conditions.

In primary human microglia, R-BHB upregulates certain inflammatory genes both with and without LPS (Supplemental Figure 4B and C, “Up by R-BHB”). However, when exposed to LPS stimulation, R-BHB had a downregulating effect on numerous genes linked to inflammation, including “TNFA Signaling via NFkB,” “Inflammatory Response,” and “Interferon Gamma Response” (Supplemental Figure 4B and C, “Down by R-BHB”). These pathways suggest that R-BHB has a modest pro-inflammatory effect without LPS while having a potent anti-inflammatory impact when microglia is activated by LPS.

In Figure 3, our in vitro experiments used both acid (endogenous product) and salt forms of BHB. However, only the acid forms were effective in reducing inflammatory molecules (Figure 3B). The introduction of acid forms of R-and S-BHB led to a small reduction in pH in the culture media to the low end of the normal range of plasma pH (7.35-7.45) (Supplemental Figure 7A). Physiologically, the production of ketone bodies (primarily acid form of R-BHB) by a KD induces metabolic acidosis that is regulated under normal conditions (but not in, for example, dysregulated diabetic ketoacidosis)^49^. However, a recent study shows a KD induces intracerebral acidosis (from pH 7.2 to 6.9) which has anticonvulsant effects in a rat model of infantile spasms^50^. R-BHB at 5 mM (but not 1 mM) reduces inflammation in primary microglia and the 5 mM slightly decreases the pH in the culture medium (Supplemental Figure 7A and B). In contrast, salt forms of R-and S-BHB do not alter the pH (Supplemental Figure 7A) or impact expressions of inflammatory molecules (Figure 3B). Minor variations in the culture media’s pH may potentially affect inflammation modulation. In a study on microbiota, Ang et al. propose that the acidic form of BHB inhibits the growth of Bifidobacterium in vitro by a pH-dependent mechanism^51^. Further investigations are necessary to clarify the pH-dependent roles of BHB acids in the transcriptome, specifically regarding inflammation.

The primary role of BHB is to provide energy to cells and tissues^2^. Macrophages do not oxidize BHB^52^. However, neurons^53,54^, CD4^+^ T cells^34,35^, CD8^+^ T cells^34,36^, and microglia^41^ do, results in the formation of ATP and intermediate metabolites, such as acetyl-CoA and succinate. Although S-BHB is oxidized much more slowly than R-BHB^2^, both forms of BHB acid reduced inflammation to a similar degree (Figure 3B), suggesting both R-and S-BHB appear to function as signaling molecules rather than energy metabolites for achieving anti-inflammatory effects.

BHB is a known inhibitor of class I HDACs^6^, and BHB regulates the activities of neuronal and stem cells via inhibiting HDACs^55–57^. HDAC inhibitors have been shown to possess anti-neuroinflammatory functions^58^. Our study found that the HDAC inhibitor butyrate (Bu and Na-Bu) significantly decreased inflammation in primary microglia (Figure 3B). Additionally, the RNA-seq analysis presented in Figures 2B and C indicates that R-BHB inhibits “G2-M Checkpoint” genes in all cells, which is consistent with the overall effect of HDAC inhibitors, which induce cell cycle arrest and have anti-cancer properties^59^. It indicates that BHBs could act as an HDAC inhibitor in microglia. Huang et al. support this hypothesis by illustrating that BHB causes ramification of microglia through HDAC inhibition^42^. However, we are not able to validate the hypothesis in either mouse primary microglia or other cell types utilized during this study. In contrast to butyrate (Bu and Na-Bu), BHBs did not induce histone acetylation (H3K9ac and total Kac around 17kD (molecular weight of histone H3)) in microglia (Figure 3C). Other studies have similarly shown that BHBs may not elevate H3K9ac levels in certain cell types^7,60^, supporting the notion that overall BHB effects on histone acetylation could be cell-specific, perhaps due to compensation of HDAC inhibition by other competing mechanisms. One possibility is that BHB may impact the equilibrium of cytoplasmic and mitochondrial nicotinamide adenine dinucleotide (NAD^+^) while activating sirtuins, NAD^+^-dependent histone deacetylases, and promoting histone deacetylation^2,61^, Further investigations may be required to understand the relationship between BHB and NAD^+^ metabolism in the brain.

The induction of H3K9bhb levels by BHB is critical for CD8^+^ T cell cells^62^ and small intestinal epithelial^63^ functions. We noted slight induction of H3K9bhb levels by BHBs in microglia, but butyrate (Bu and Na-Bu) is a more potent inducer (Figure 3C), which was already reported in other cell types^60^. We observed a significant increase in the expression of total protein Kbhb with R-BHB and Na-R-BHB treatments (this antibody specifically detects the R form of Kbhb) but not with Bu or Na-Bu treatments (Figure 3C). Contrary to IMG cells (Supplemental Figure 10C), BV-2 cells (Supplemental Figure 11C), and BMDMs (Supplemental Figure 12C), we were unable to detect Kbhb bands at approximately 17 kD in primary mouse microglia (Figure 3C) and astrocytes (Supplemental Figure 9B) by R-BHB and Na-R-BHB. As reported recently, the H3K9bhb antibody can potentially recognize non-specific modifications of histones^64^. Koronowski et al. showed that after four weeks of a KD, there is an induction of total protein Kbhb in the liver, but not in the brain^65^. While a one-week KD leads to a 15-20-fold increase in R-BHB levels in the brain^37^, there were no noted changes in total protein Kac or Kbhb levels after one week of the diet (Supplemental Figure 12C). The findings suggest that a KD could have unique effects on metabolic pathways within specific tissues, with Kbhb potentially regulated in a dissimilar manner between the brain and other tissues, such as the liver.

We did not observe a decrease in NLRP3 inflammasome activation by BHBs across all cell types, despite the control inhibitor MCC950 performing consistently. Interestingly, acid forms of BHBs even amplified nigericin-induced inflammasome activation in primary microglia (Supplemental Figure 7B, C). Following the previous experiments^4^, we administered these compounds with secondary activators ATP/nigericin (Supplemental Figure 7A). A study utilizing mouse primary microglia indicated that BHB impedes the activation of NLRP3 inflammasome by ATP or monosodium urate (MSU), but not by synuclein^66^. A recent report showed BHB reduces amyloid beta oligomer (AβO)-induced NLRP3 inflammasome in human induced pluripotent stem cell (iPSC)-derived microglia^67^. In an *in vivo* model, BHB administration inhibits the activation of NLRP3 inflammasome in the cortices of 5XFAD AD mice^43^. Investigating the mechanisms by which BHB regulates NLRP3 inflammasome in the brain is necessary.

BHB is known to be a GPCR Hcar2 agonist, and in an ischemic stroke model, Rahman et al. demonstrated that the activation of Hcar2 by BHBs in brain-infiltrated monocytes/macrophages is neuroprotective^44^. Activation of Hcar2 in microglia appears to be neuroprotective in Alzheimer’s disease^68^. Furthermore, a recent study suggested BHB suppresses the growth of intestinal epithelium and colorectal cancer via this receptor^69^. The receptor’s role in neuroinflammation, however, is unclear. In primary microglia, LPS induced an over 100-fold increase in Hcar2 expression. Reducing the gene expression by 50% did not affect inflammation in primary microglia. (Supplemental Figure 7D). In vascular cells (smooth muscle cells and endothelial cells), BHB binds directly to hnRNP A1, upregulating Oct4, an Octamer-binding transcriptional factor, and preventing senescence. Although Oct4 is not expressed in microglia (not shown), BHB could decrease inflammation in a Hnrnpa1-dependent manner. However, a 50% decrease in Hnrnpa1 expression in microglia did not alter the effects of BHB on inflammation (Supplemental Figure 7D). In future studies, it may be necessary to explore more rigorous methods to investigate the anti-inflammatory mechanism of BHB acid, such as utilizing knockout mouse models and screening for BHB binding proteins in microglia.

To our knowledge, no publications have investigated the *in vitro* transcriptomic effects of BHB on both neuronal and glial brain cells. Using RNA-seq, Ruppert et al. analyzed the transcriptional changes induced by BHB and butyrate in various primary cell types, such as BMDMs^70^. During the experiments, a 5 mM salt form of R-BHB (Na-R-BHB) was used for six hours without any stimulation, such as LPS. From their findings, the authors concluded that Na-R-BHB had minimal impact on gene expression in macrophages. The transcriptional effects of BHB depend on various factors, including cell type, incubation time, concentration, and formulation. In our study, we observed the anti-inflammatory effects of R-BHB (acid) in BMDMs (Supplemental Figure 11A), suggesting that the acid form of R-and S-BHB may exhibit anti-inflammatory effects not only in microglia but also in other immune cell types, particularly macrophages.

In Figure 4, we utilized bulk RNA-seq to analyze the brains of mice fed either a CD or a KD. For the one-week cohort, we employed a one-week ad libitum KD feeding regimen in 12-month-old mice. As previously noted, high levels of R-BHB is expected in the brains of KD-fed mice during this feeding regimen^37^. During the one-year cohort study, we implemented a cyclic KD feeding plan every other week from 12 to 26 months of age. Samples were collected during the CD-fed period. It should be noted that blood R-BHB levels of cyclic KD-fed mice at the CD-fed period were identical to those of ad libitum CD-fed mice^10^. The levels of R-BHB in brain (and liver) tissue from mice on a cyclic KD diet should be equivalent to those of mice on a standard CD diet.

The results demonstrate that a one-week KD increased various signaling pathways, including “Hedgehog Signaling”, “Notch Signaling”, and “Inflammatory Response” (Supplemental Figure 12B). *In vitro* experiments indicated that R-BHB (without LPS) increased “Inflammatory Response” but did not affect “Hedgehog Signaling” and “Notch Signaling” in all cell types (Figure 2B). However, BHB enhances Notch signaling in intestinal stem cells^57^ and the “G2-M Checkpoint” signature observed *in vitro* (Figure 2B and C), did not change *in vivo* after the one-week KD treatment. These findings suggest that the effects of the KD on the brain are not solely due to the induction of R-BHB, but may also involve other mechanisms, such as the absence of carbohydrates. Because the effects of one-year cyclic KD should not depend on the direct effects of R-BHB on brain cells, the mechanisms by which the feeding regimen reduces age-induced chronic inflammation are not simple. For instance, there may be cell type specificity and dependency on R-BHB. One possibility is that chronic exposure to R-BHB in the brain leads to epigenetic changes in the protein Kbhb in a cell-specific manner, but we did not observe any induction of Kbhb in the brain after a one-week KD (Supplemental Figure 13C).

One potential factor to consider is the link between KD and systemic inflammatory effects. Goldberg et al. utilized a single-cell RNA-seq to demonstrate that a one-week KD can modify the immune population in the adipose tissues^31^. The one-week KD was shown to reduce the population of pro-inflammatory macrophages and increase metabolically protective γδ T cells in the adipose tissues. Conversely, a 2-3-month ad libitum KD causes adipose inflammation by increasing macrophages and decreasing γδ T cells. In our study, a one-week KD did not result in any notable changes in inflammation in the liver and brain. However, a one-year cyclic KD significantly decreased chronic inflammation in both tissues (Figures 1B and 4B). These findings suggest that extended KD therapy with sufficient breaks (every other week) may decrease persistent inflammation via γδ T cell stimulation. In a separate study, Goldberg and colleagues indicate that metabolic adaptation through KD, rather than induction of BHB, is necessary for the expansion of γδ T cells^71^. We demonstrate that a cyclic KD can reverse the age-related phenotype of blood T cells (Supplemental Figure 2B). This indicates that the systemic anti-inflammatory effects of KD may have a role to play in the brain.

The changes in gene expression triggered by R-BHB in microglia are mostly in line with those caused by KD treatments in the brain (Supplemental Figure 15C-E). This indicates that the anti-inflammatory effects of KDs are probably rooted primarily in microglia. At a single-cell resolution analysis of the mouse brain, the upregulated genes with age are generally associated with inflammatory and immune responses in glia, particularly microglia and astrocytes^40^. However, our study does not explore the cell-and region-specific functions of KDs in the brain *in vivo*, and there have been no investigations into the cell-specific transcriptional changes in the brain following KD treatments in aging mammal models. To achieve this, a single-cell RNA-seq should be employed in precise brain regions, such as the cortex, hippocampus, and hypothalamus.

Using only young mice, two studies have investigated the cell-specific role of KD using transcriptional or proteomic approaches^72,73^. Koppel et al. utilized bulk RNA-seq to analyze neuron-or astrocyte-enriched fractions treated with a KD. The study utilized 16-week-old mice on a three-month ad libitum KD and found differential effects on neurons and astrocytes^72^. The study indicates activation of inflammation-associated signaling in astrocytes, but not in neurons. It appears that a 3-month ad libitum KD is sufficient to induce inflammation in both the brain and adipose tissue^31^. Düking et al. employed cell type-specific proteomics to analyze KD-treated brain cells, encompassing neurons, astrocytes, and microglia^73^. The study utilized 14-day-old pups and a 4-week ad libitum KD, demonstrating that the KD increased proteins relating to ketolysis (such as OXCT1/SCOT), the TCA cycle, and OXPHOS in astrocytes and neurons but not microglia. In our study, we did not observe the induction of Oxct1 by KDs in the brain. However, after a one-week KD, genes related to fatty acid oxidation (Cpt1a) and ketogenesis (Hmgcs2 and Hmgcl) were induced in the brain (Supplemental Figure 13E) as well as the liver (Supplemental Figure 1D). As we were unable to detect the ketogenic gene HMGCS2 in all brain cells *in vitro* (Supplemental Figure 4E), a clear understanding of the role of fatty acid oxidation and ketogenesis in the KD-fed brain remains elusive. Recent research has demonstrated that fatty acid oxidation in adult mouse astrocytes is crucial for cognitive function^74^. In Drosophila, glial cells provide energy to neurons through ketogenesis, which is necessary for memory and survival during periods of starvation or when glial glycolysis is absent^75,76^. Further studies are necessary to investigate the role of ketogenesis in the mammalian brain.

In summary, our study demonstrates that the endogenous metabolite R-BHB reduces LPS-induced inflammation in microglia (Figure 2). This function is likely independent of energy supply, HDAC inhibition, or Hcar2 activation (Figure 3). Additionally, we observe that a cyclic KD reduces age-induced chronic inflammation in the brain (Figure 4), as well as in the liver (Figure 1). As previously noted, our research demonstrated an improvement in lifespan and memory function in aging mice with a cyclic KD^10^. When conducting translational studies on KD feeding, it may be important to prevent obesity and metabolic maladaptation, and to take sufficient breaks during a long-term dietary intervention. Our study shows that sporadic KD feeding could serve as a feasible dietary intervention for human aging by preventing chronic inflammation, a hallmark of aging^33,77^.

## Acknowledgments

This work was supported by the NIH grants R01AG067333 (J.C.N.), T32AG000266 (M.N), T32AG052374 (S.S.M. and B.E.), Buck intramural funds (J.C.N.), and University of Southern California Provost Fellowship Funding (S.S.M.). Both IMG and BV-2 cells were kindly provided by the Andersen Lab. Some illustrations were created with BioRender.com.

## Authorship Contributions

M.N. conceived the studies, designed the experiments, analyzed experiments, and wrote the manuscript. N.F.M. and S.S.M. analyzed the RNA-seq data. T.Y.G., B.E., and C.G.A. helped with experiments. L.E. and D.F. provided important insights and suggestions to the manuscript. E.V. supervised in vivo cohorts. J.C.N. supervised the whole study and co-wrote the manuscript.

## Data Availability Statement

Raw data sets for RNA-seq are deposited in GEO.

