## Supplemental for "A ketogenic diet reduces age-induced chronic neuroinflammation in mice"

**Supplemental Figure Legends**

**Supplemental Figure 1. Related to Figure 1**

(A) Volcano plot of gene expressions (KD vs. CD) of 26-month liver on diets for 14 months. Red (upregulated) and blue (downregulated) dots represent genes with a P value less than 0.02. (B) GSEA (GO) analysis in the liver. (C) GSEA (KEGG) analysis in the liver. (D) Heatmap of gene expression (KD vs. CD) of one-week on diets in the 12-month-old liver (n=5 (CD) and 7 (KD) mice per group).

**Supplemental Figure 2. Related to Figure 1**

(A) Experimental timeline with mouse numbers. Chow is the standard vivarium chow. (B) $CD4^+$ cells, $CD8^+$ cells, and $CD8^+/CD4^+$ ratio in the blood. (C) $Foxp3^+$ cells in the blood. (D) $PD\text{-}1^+/CD4^+$ and $PD\text{-}1^+/CD8^+$ cells in the blood. All data are presented as mean ± SD. One-way ANOVA with Dunnet's correction for multiple comparisons.

**Supplemental Figure 3. Related to Figure 2**

(A) mRNA expression in human primary microglia (n=2 per each group). All data are presented as mean ± SD. (B-D) Volcano plots of gene expressions. Red (upregulated) and blue (downregulated) dots represent genes with a P value less than 0.02. (E) GSEA (hallmark) analysis (+ LPS vs. − LPS) in human primary neurons/astrocytes/microglia.

**Supplemental Figure 4. Related to Figure 2**

(A) Heatmap of "G2-M Checkpoint" genes. (B) Heatmap of "TNF Alpha Signaling via NFkB" genes. (C) Heatmap of "Inflammatory Response" genes. (D) Heatmap of "Interferon Gamma Response" genes. (E) Heatmap of genes related to ketone metabolism and monocarboxylate transporters (MCTs).

**Supplemental Figure 5. Related to Figure 2**

(A-C) GSEA (GO) analysis in human primary neurons (A), astrocytes (B), and microglia (C).

**Supplemental Figure 6. Related to Figure 2**

(A-C) GSEA (KEGG) analysis in human primary neurons (A), astrocytes (B), and microglia (C).

**Supplemental Figure 7. Related to Figure 3**

(A) Compound structures and the pH in the compounds' culture media. (B-D) mRNA expression in mouse primary microglia (n=3 per group). All data are presented as mean ± SD. One-way ANOVA with Dunnet's correction for multiple comparisons. (B, C) Compare the mean of each sample with the mean of 0 mM. All data are representative of two independent experiments.

**Supplemental Figure 8. Related to Figure 3**

(A) Experimental timeline. (B) ELISA analysis of IL-1β secretion after NLRP3 inflammasome activation in mouse primary microglia (n=3 per group). All data are presented as mean ± SD. One-way ANOVA with Dunnet's correction for multiple comparisons. Compare the mean of each sample with the mean of Ctrl (+ ATP or nigericin). (C) WB analysis of IL-1β secretion after NLRP3 inflammasome activation in mouse primary microglia (n=1-2 per group). All data are representative of two independent experiments.

**Supplemental Figure 9. Related to Figure 3**

(A) mRNA expression in primary astrocytes (n=3 per group). All data are presented as mean ± SD. One-way ANOVA with Dunnet's correction for multiple comparisons. Compare the mean of each sample with the mean of Ctrl. (B) Protein expression in mouse primary astrocytes (n=2 per group). All data are representative of two independent experiments.

**Supplemental Figure 10. Related to Figure 3**

(A) mRNA expression in IMG mouse microglia cell line (n=2 per group). All data are presented as mean ± SD. One-way ANOVA with Dunnet's correction for multiple comparisons. Compare the mean of each sample with the mean of Ctrl. (B) ELISA analysis of IL-1β secretion after NLRP3 inflammasome activation in IMG cells (n=3 per group). One-way ANOVA with Dunnet's correction for multiple comparisons. Compare the mean of each sample with the mean of Ctrl (+ ATP or nigericin). (C) Protein expression in IMG cells (n=2 per group). All data are representative of two independent experiments.

**Supplemental Figure 11. Related to Figure 3**

(A) mRNA expression in BV-2 mouse microglia cell line (n=2 per group). All data are presented as mean ± SD. One-way ANOVA with Dunnet's correction for multiple comparisons. Compare the mean of each sample with the mean of Ctrl. (B) ELISA analysis of IL-1β secretion after NLRP3 inflammasome activation in BV-2 cells (n=3 per group). All data are presented as mean ± SD. One-way ANOVA with Dunnet's correction for multiple comparisons. Compare the mean of each sample with the mean of Ctrl (+ ATP or nigericin). (C) Protein expression in BV-2 cells (n=2 per group). All data are representative of two independent experiments.

**Supplemental Figure 12. Related to Figure 3**

(A) mRNA expression in mouse bone marrow-derived macrophages (BMDMs) (n=2 per group). All data are presented as mean ± SD. One-way ANOVA with Dunnet's correction for multiple comparisons. Compare the mean of each sample with the mean of Ctrl. (B) ELISA analysis of IL-1β secretion after NLRP3 inflammasome activation in BMDMs (n=3 per group). All data are presented as mean ± SD. One-way ANOVA with Dunnet's correction for multiple comparisons. Compare the mean of each sample with the mean of Ctrl (+ ATP or nigericin). (C) Protein expression in BMDMs (n=2 per group). All data are representative of two independent experiments.

**Supplemental Figure 13. Related to Figure 4**

(A) Volcano plot of gene expressions in the brain. Red (upregulated) and blue (downregulated) dots represent genes with a P value less than 0.02. (B) GSEA (hallmark) analysis (KD vs. CD) of one week on diets in 12-month-old brains (n=5 (CD) and 7 (KD) mice per group). (C) Protein expressions of one-week on diets in the 12-month-old brain. (D) GSEA (hallmark) analysis of one-week on CD in 12-month-old brain (CD (12 mo)) and 26-month brain on CD for 14 months (CD (26 mo) (n=5 (CD (12 mo)) and 8 (CD (26 mo)) mice per group) (E) Heatmap of gene expressions in the brain.

**Supplemental Figure 14. Related to Figure 4**

(A-C) GSEA (GO) analysis in the brain. (D-F) GSEA (KEGG) analysis in the brain.

**Supplemental Figure 15. Related to Figure 4**

(A) RRHO analysis comparing aging (blue) with cyclic KD (brown) in the brain. (B) RRHO analysis comparing cyclic KD in the brain (blue) with liver (brown). (C) RRHO analysis comparing R-BHB (− LPS) in brain cells (blue) with one-week KD in the brain (brown). (D) RRHO analysis comparing LPS effects in brain cells (blue) with aging effects in the brain (brown). (E) RRHO analysis comparing R-BHB (+ LPS) in brain cells (blue) with cyclic KD in the brain (brown).

### Supplemental Figure 1, related to Figure 1

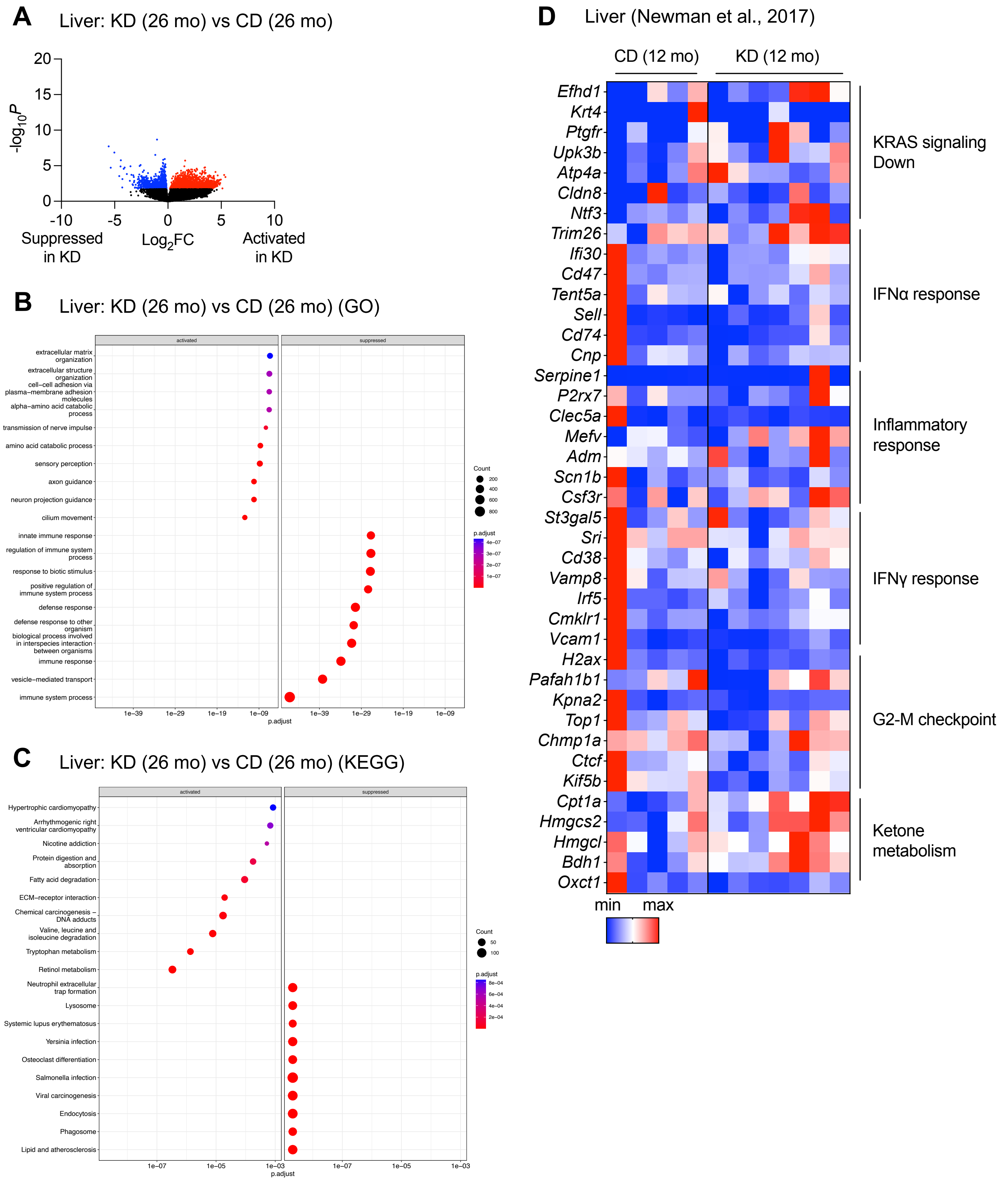

Supplemental Figure 2, related to Figure 1

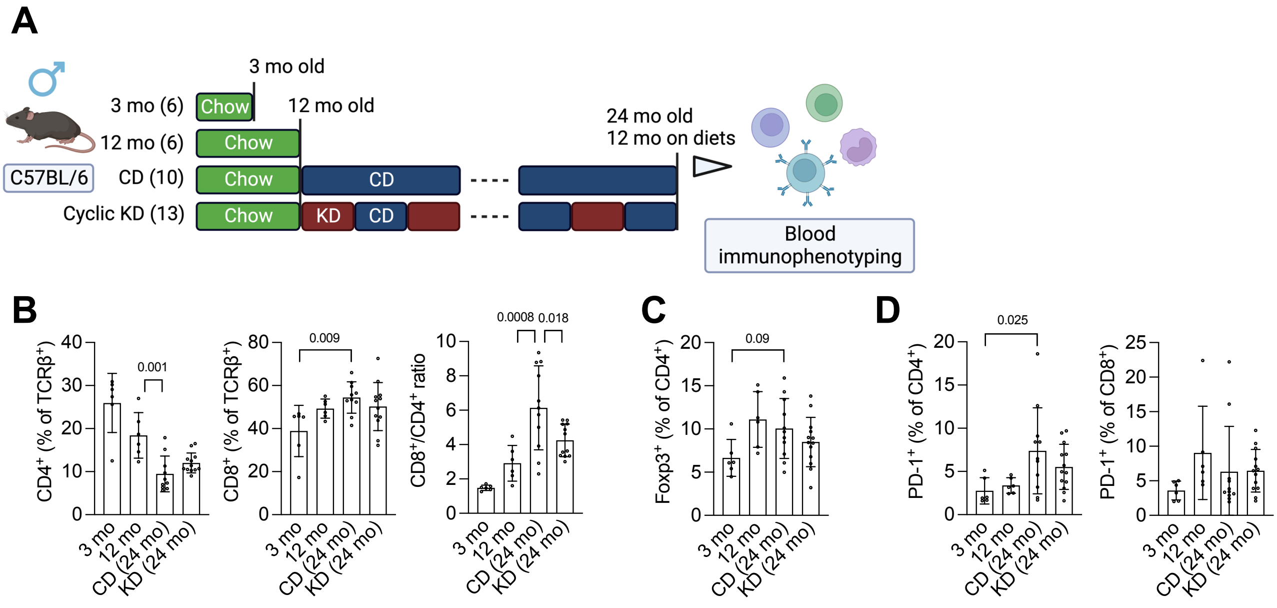

Supplemental Figure 3, related to Figure 2

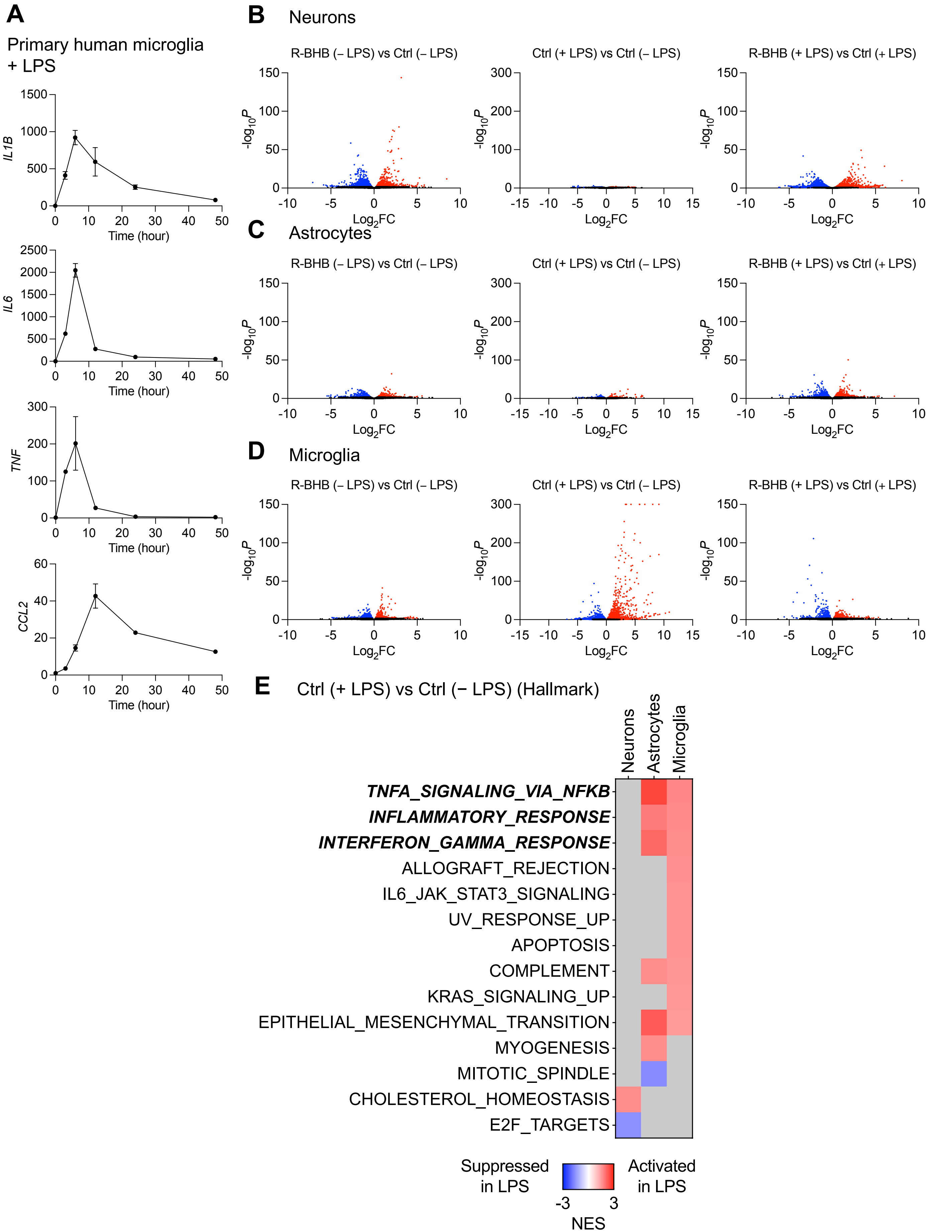

Supplemental Figure 4, related to Figure 2

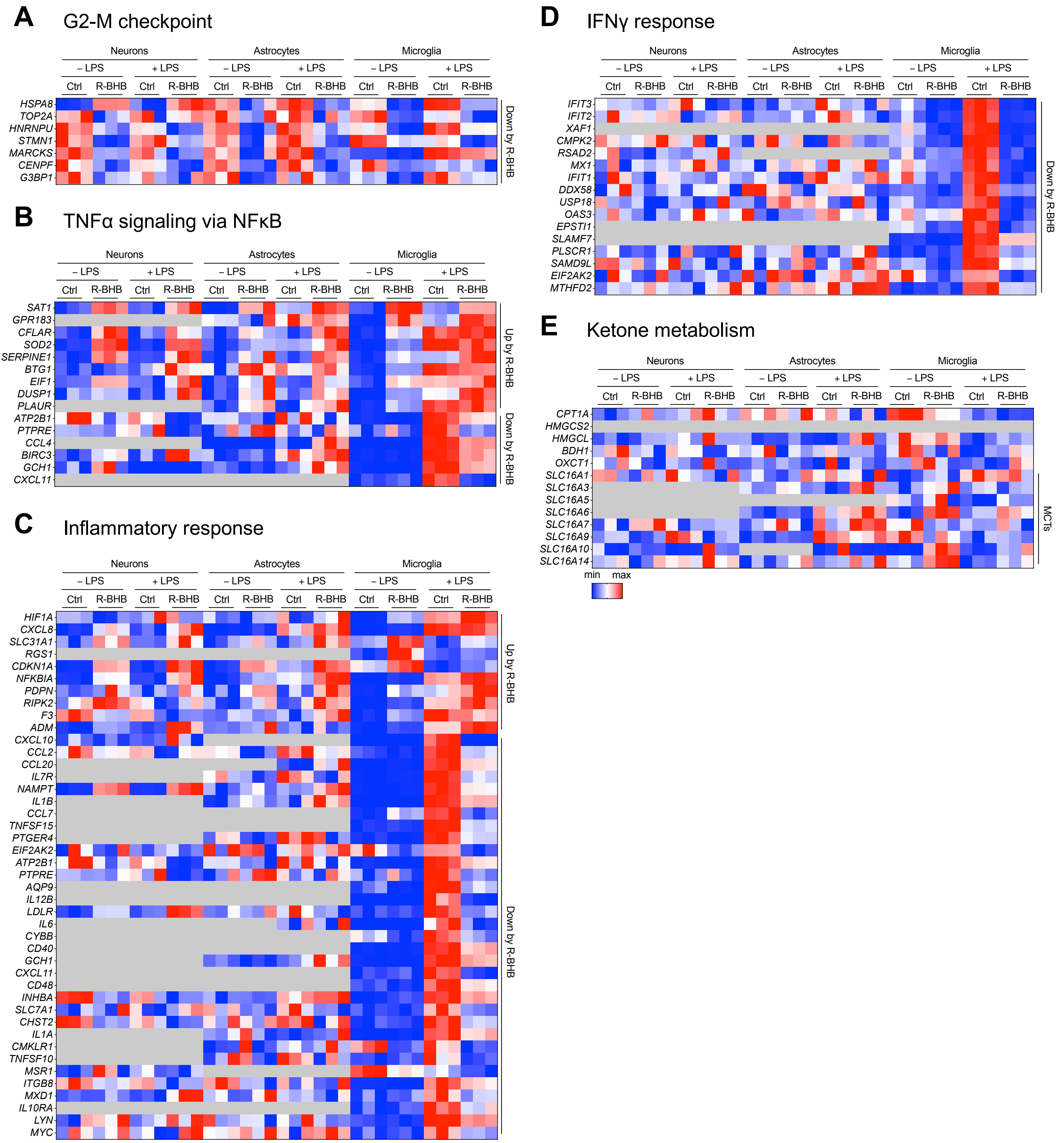

### Supplemental Figure 5, related to Figure 2

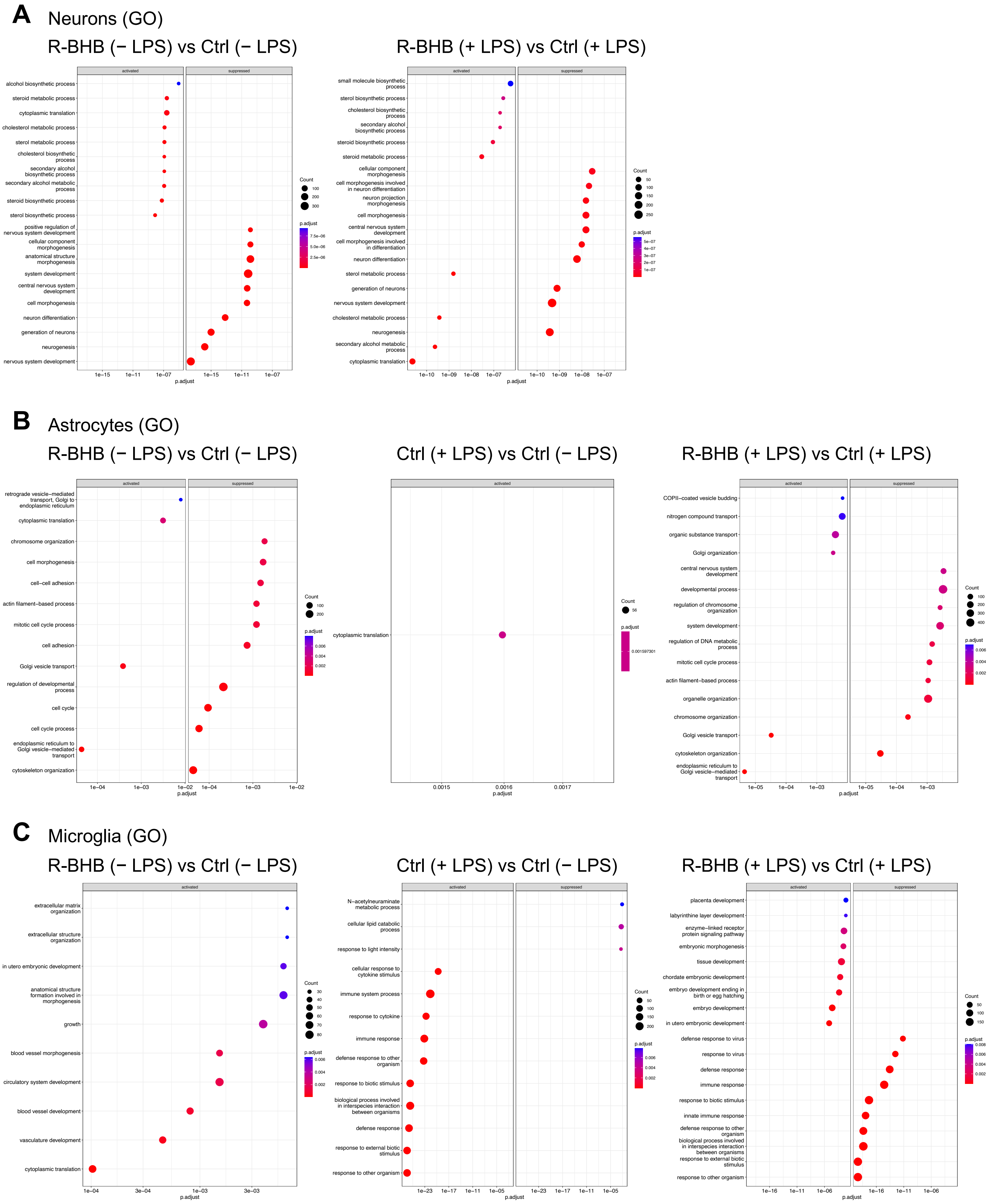

### Supplemental Figure 6, related to Figure 2

#### A Neurons (KEGG)

R-BHB (− LPS) vs Ctrl (− LPS)

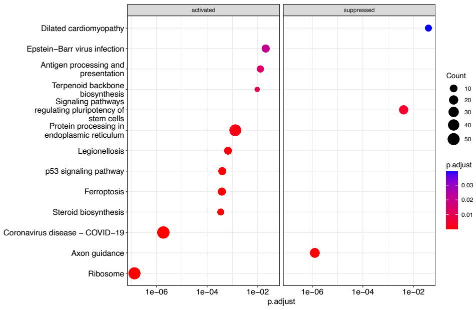


R-BHB (+ LPS) vs Ctrl (+ LPS)

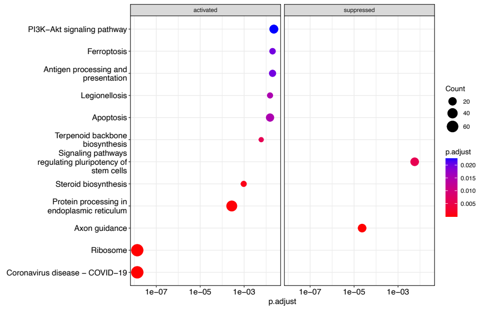


#### B Astrocytes (KEGG)

R-BHB (− LPS) vs Ctrl (− LPS)

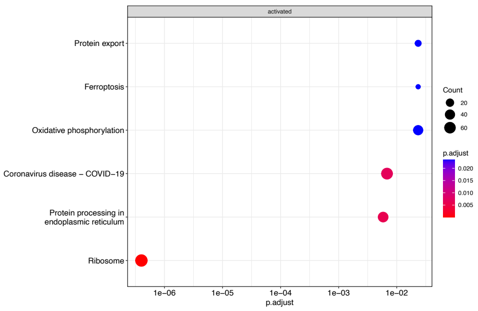


Ctrl (+ LPS) vs Ctrl (− LPS)

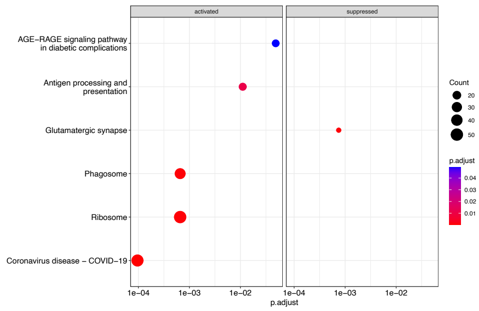


R-BHB (+ LPS) vs Ctrl (+ LPS)

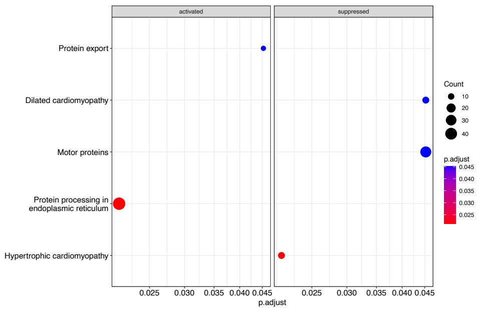


#### C Microglia (KEGG)

R-BHB (− LPS) vs Ctrl (− LPS)

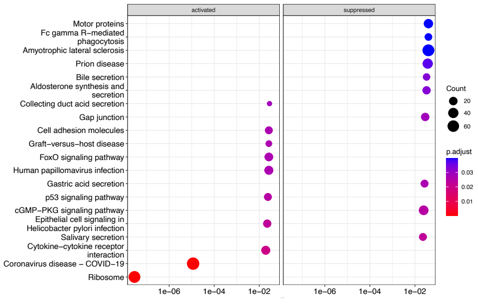


Ctrl (+ LPS) vs Ctrl (− LPS)

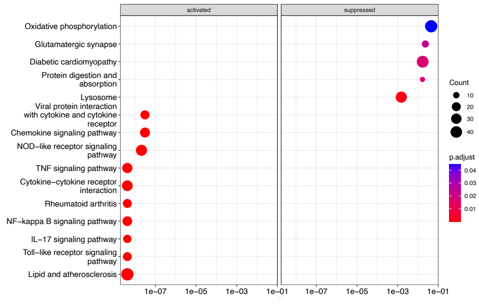


R-BHB (+ LPS) vs Ctrl (+ LPS)

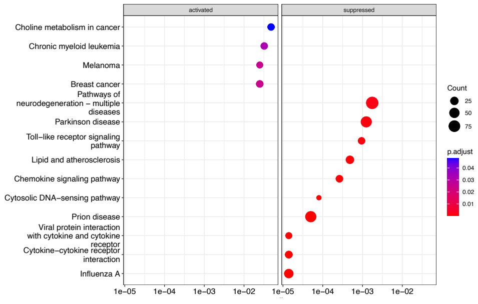

### Supplemental Figure 7, related to Figure 3

**A**

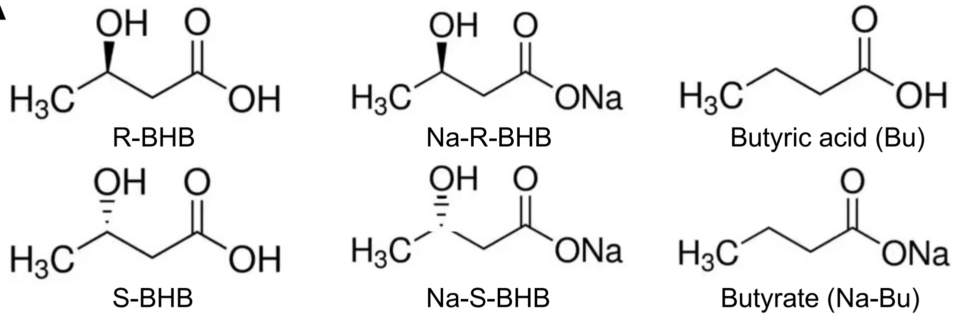


| | pH |
|---|---|
| Ctrl | 7.66 |
| R-BHB (1 mM) | 7.66 |
| R-BHB (5 mM) | 7.59 |
| R-BHB (10 mM) | 7.34 |
| S-BHB (10 mM) | 7.34 |
| Na-R-BHB (10 mM) | 7.67 |
| Na-S-BHB (10 mM) | 7.64 |
| Bu (5 mM) | 7.53 |
| Na-Bu ( 5 mM) | 7.63 |

**B** Primary microglia + R-BHB (0, 1, 5, 10 mM)

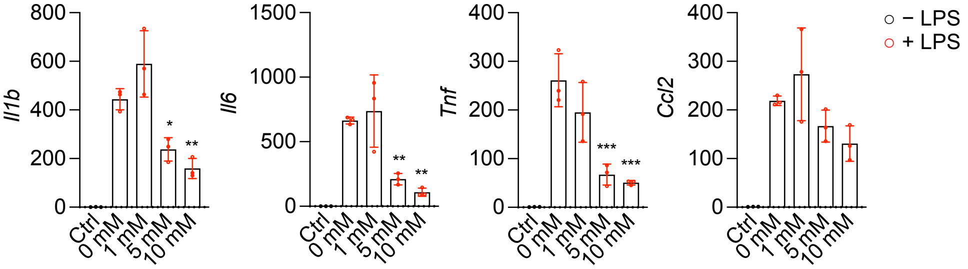


**C** Primary microglia ± R-BHB ± 10 mM NaCl

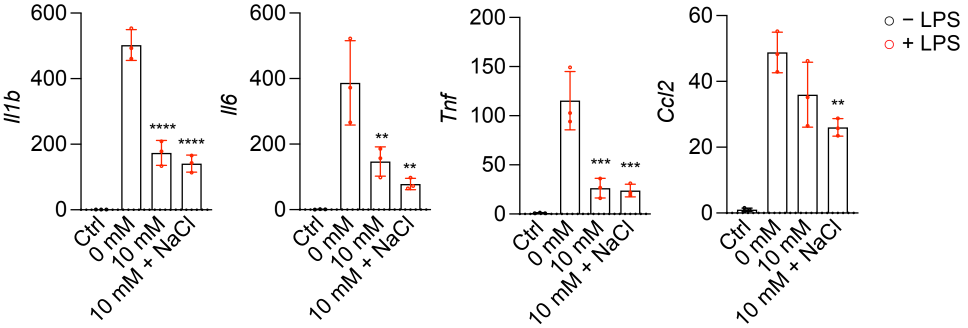


**D** Primary microglia ± R-BHB (10 mM) ± siHcar2 ± siHnrnpa1

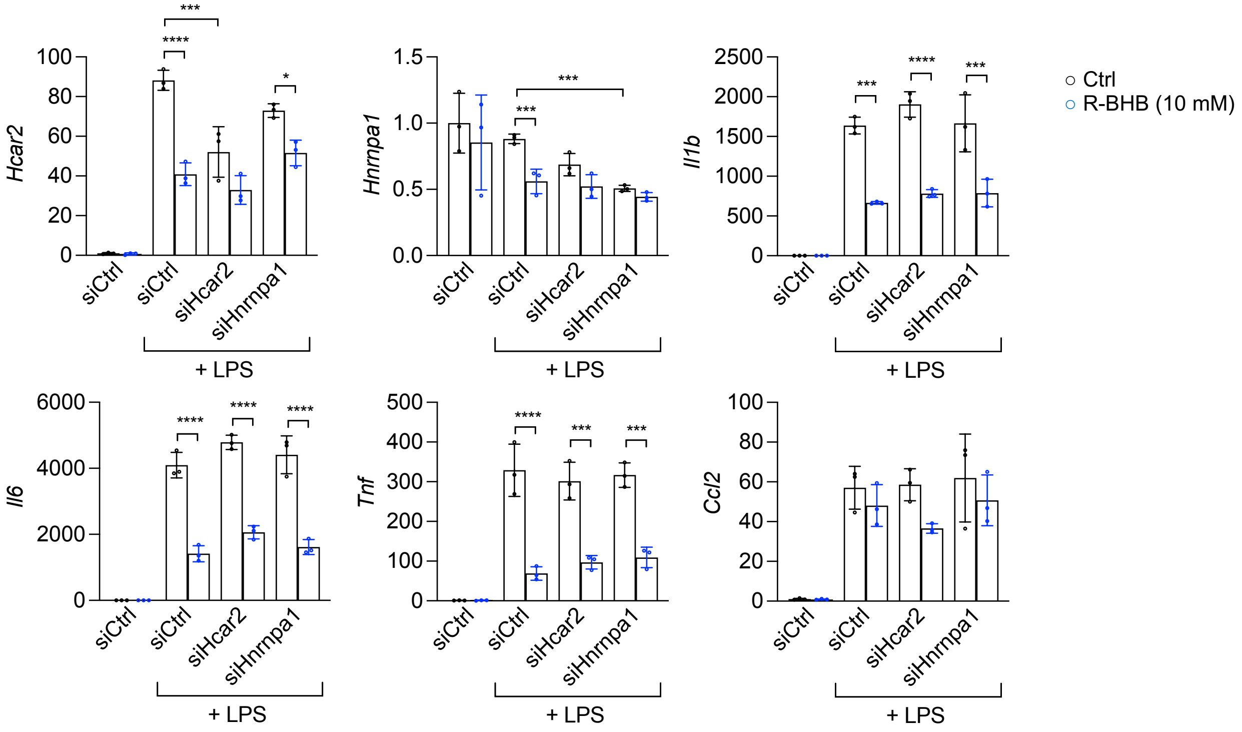

Supplemental Figure 8, related to Figure 3

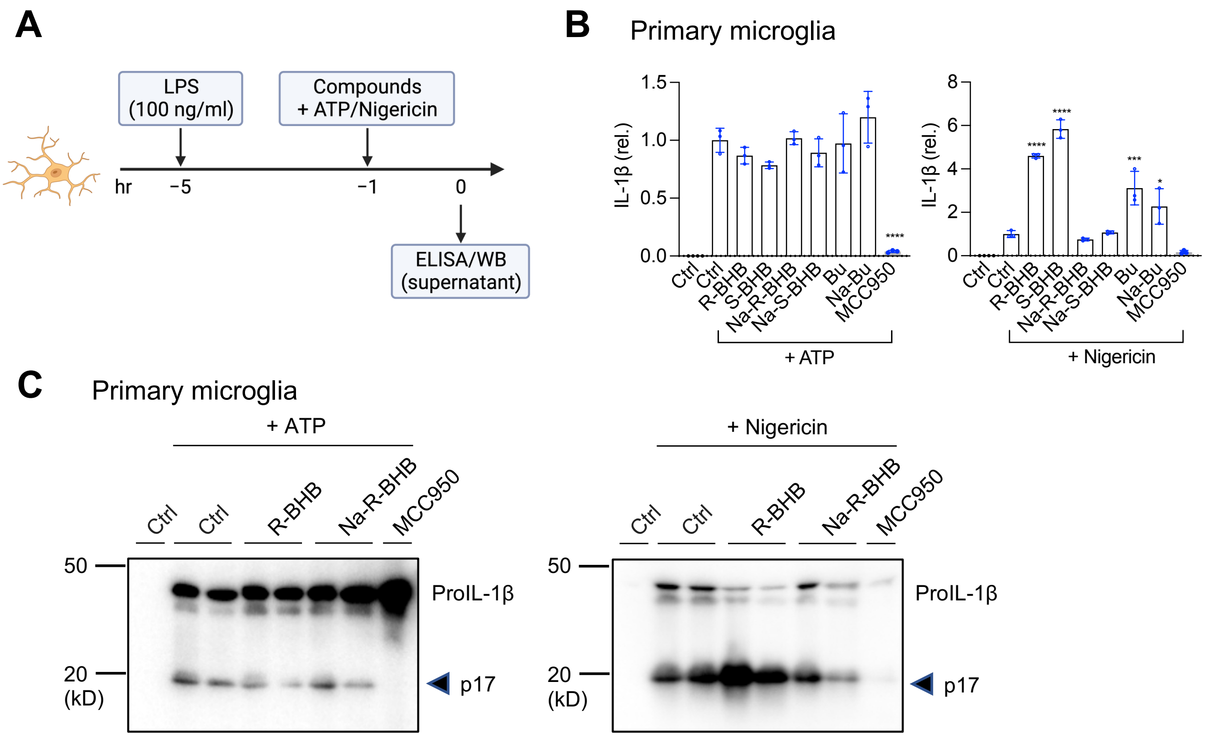

### Supplemental Figure 9, related to Figure 3

**A** Primary astrocytes

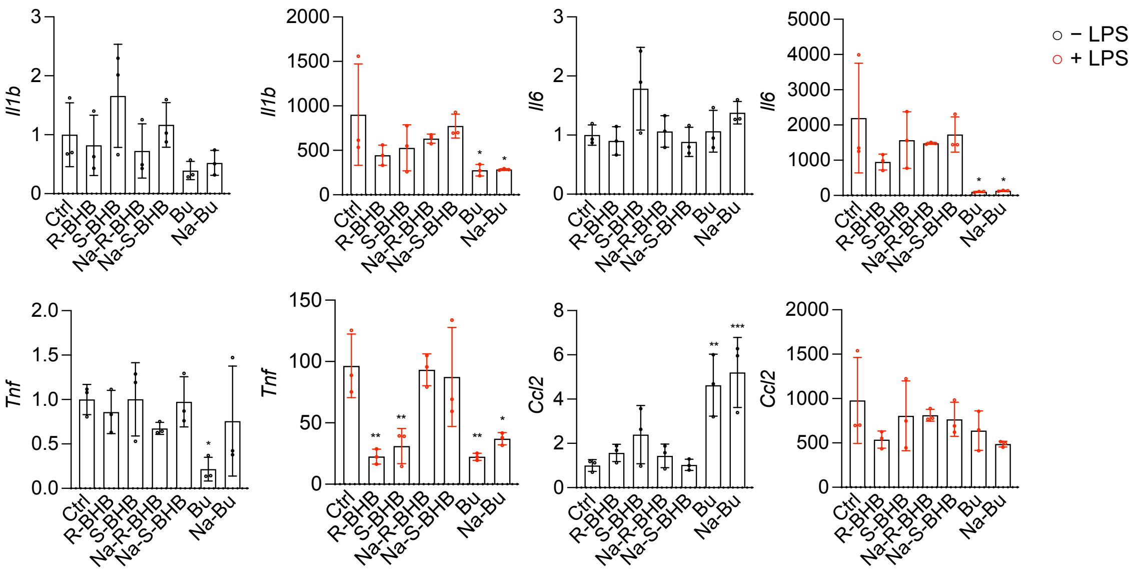


**B** Primary astrocytes

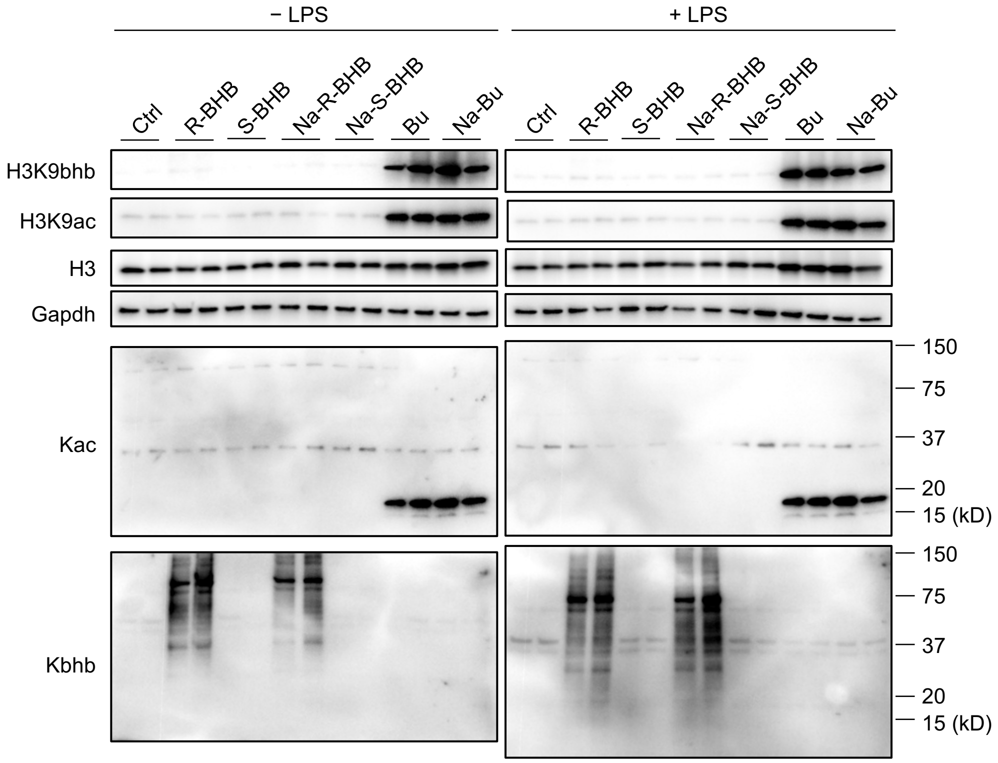

Supplemental Figure 10, related to Figure 3

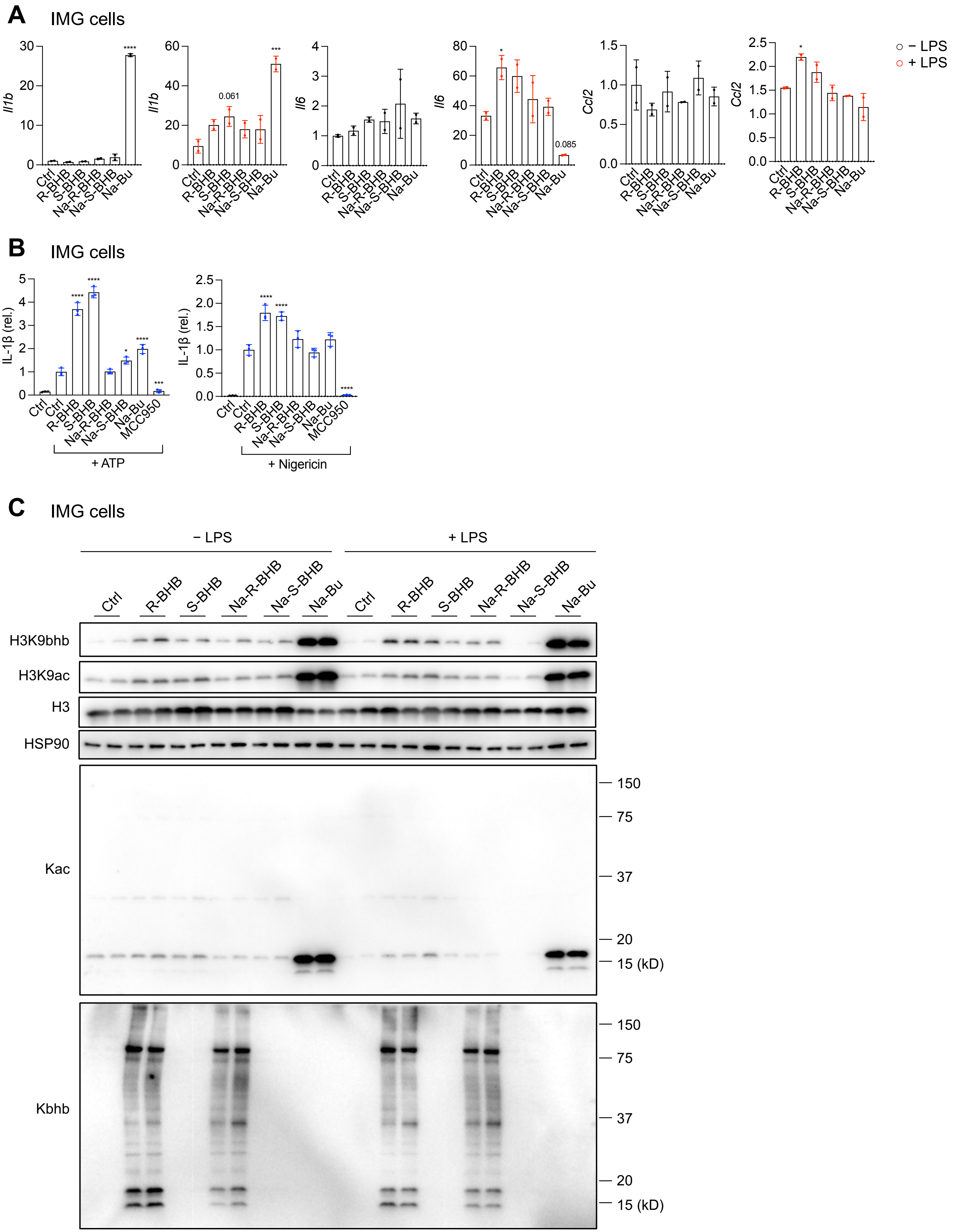

Supplemental Figure 11, related to Figure 3

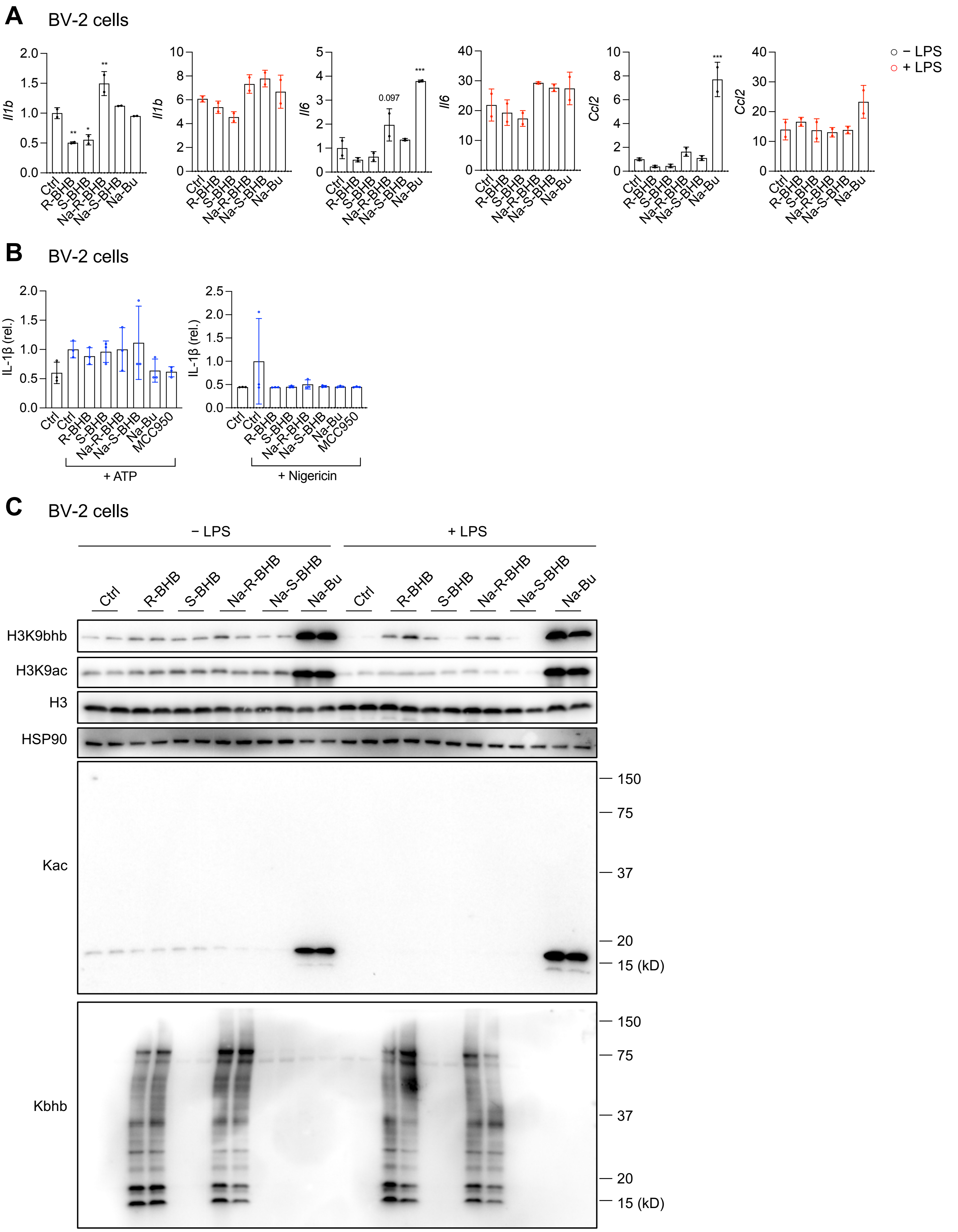

Supplemental Figure 12, related to Figure 3

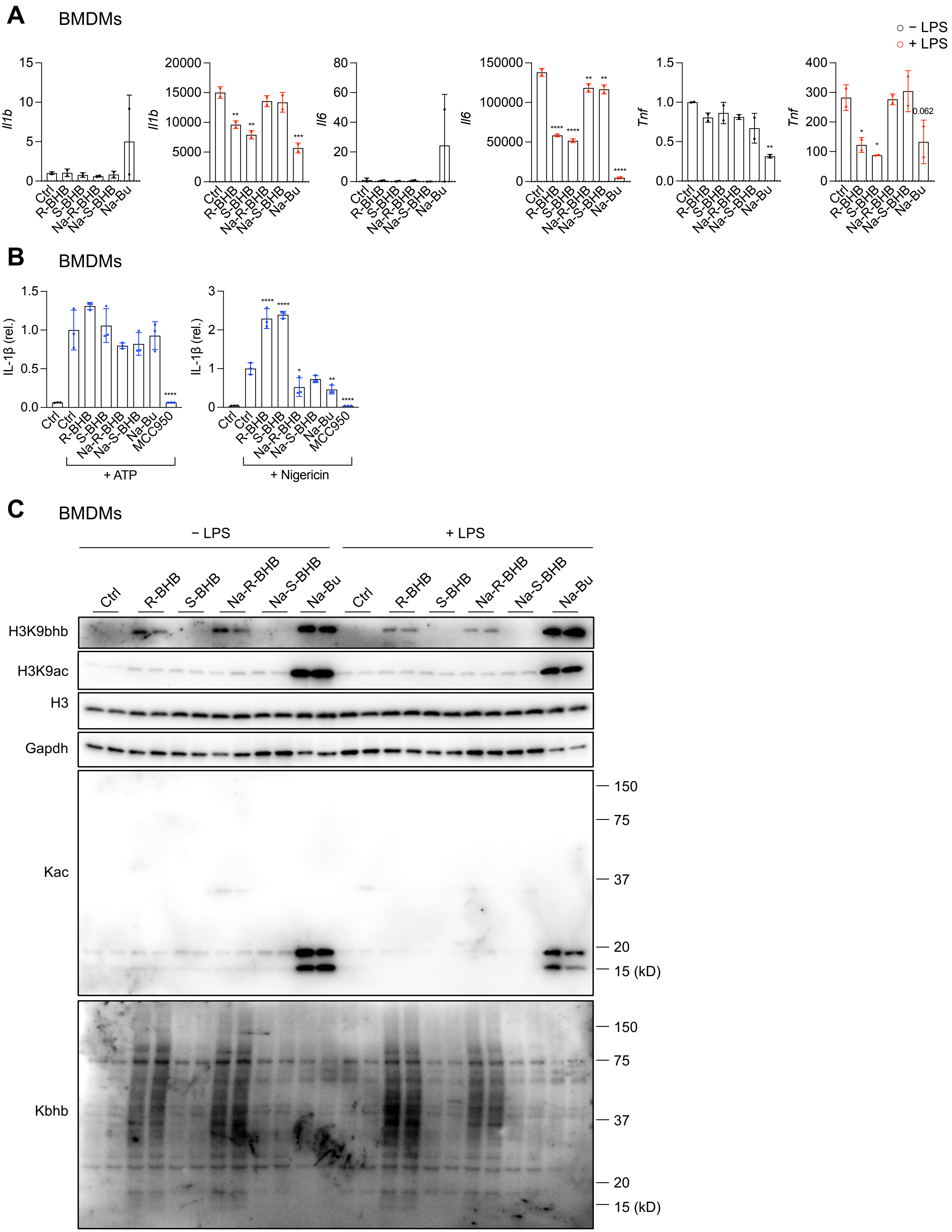

Supplemental Figure 13, related to Figure 4

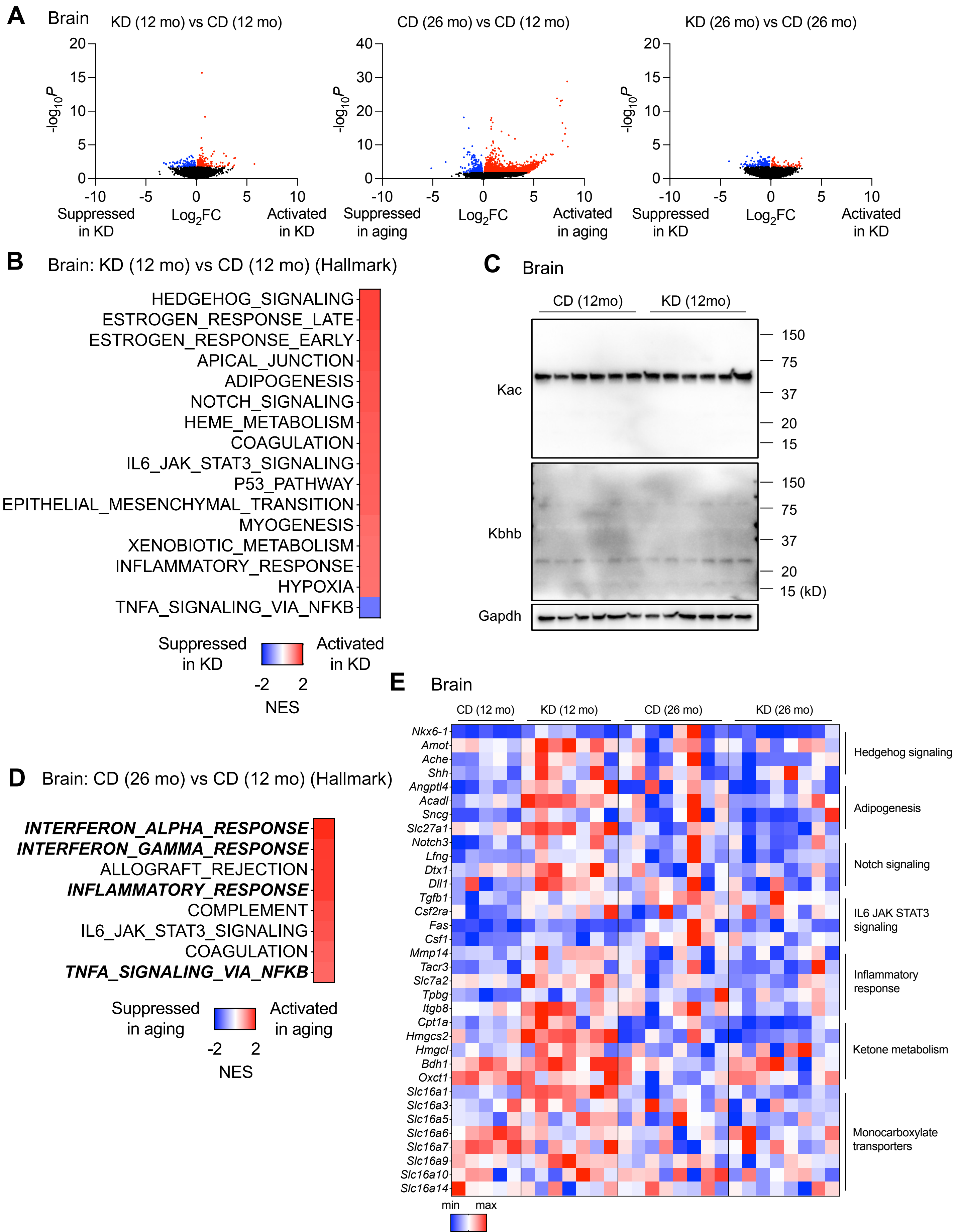

### Supplemental Figure 14, related to Figure 4

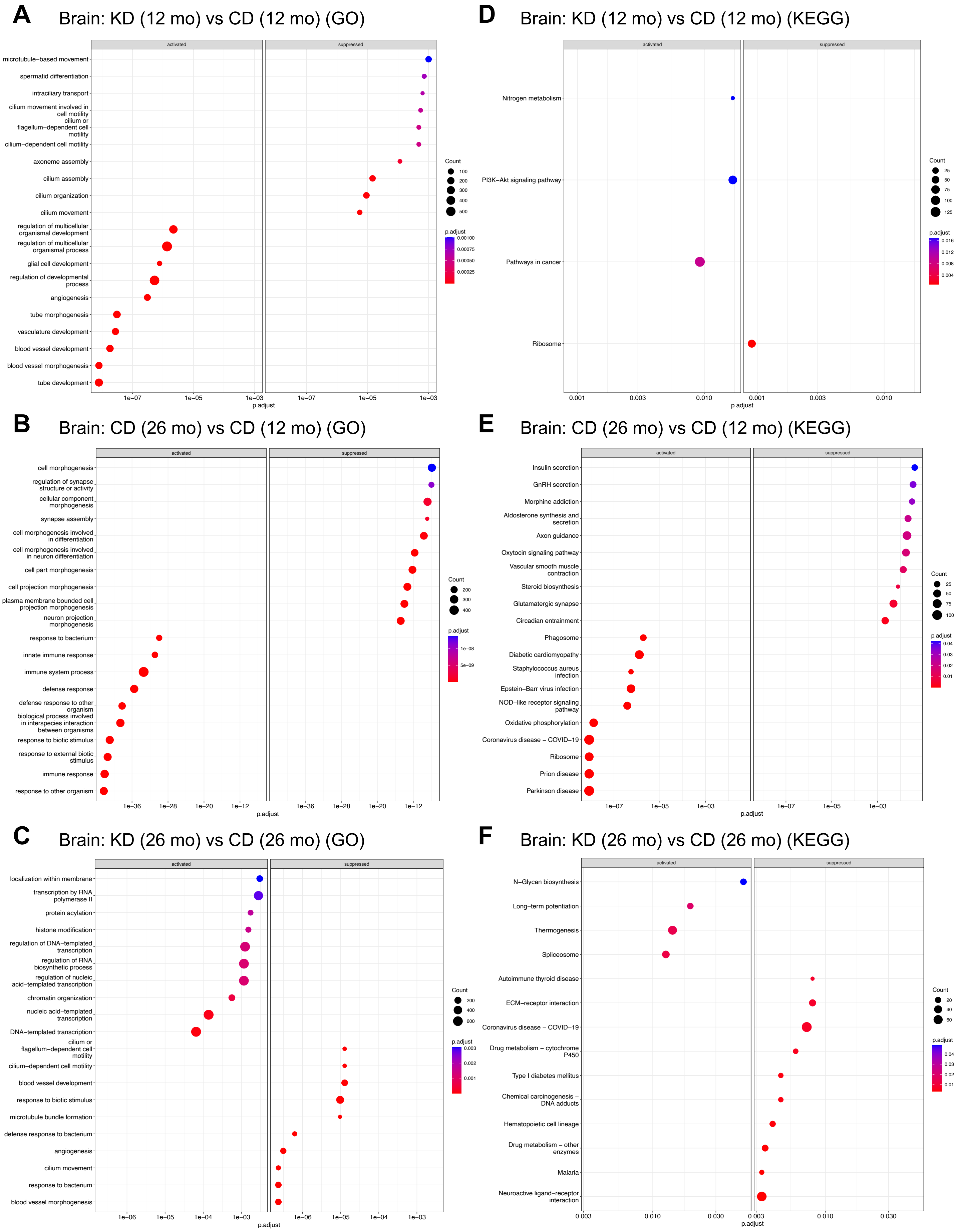

Supplemental Figure 15, related to Figure 4

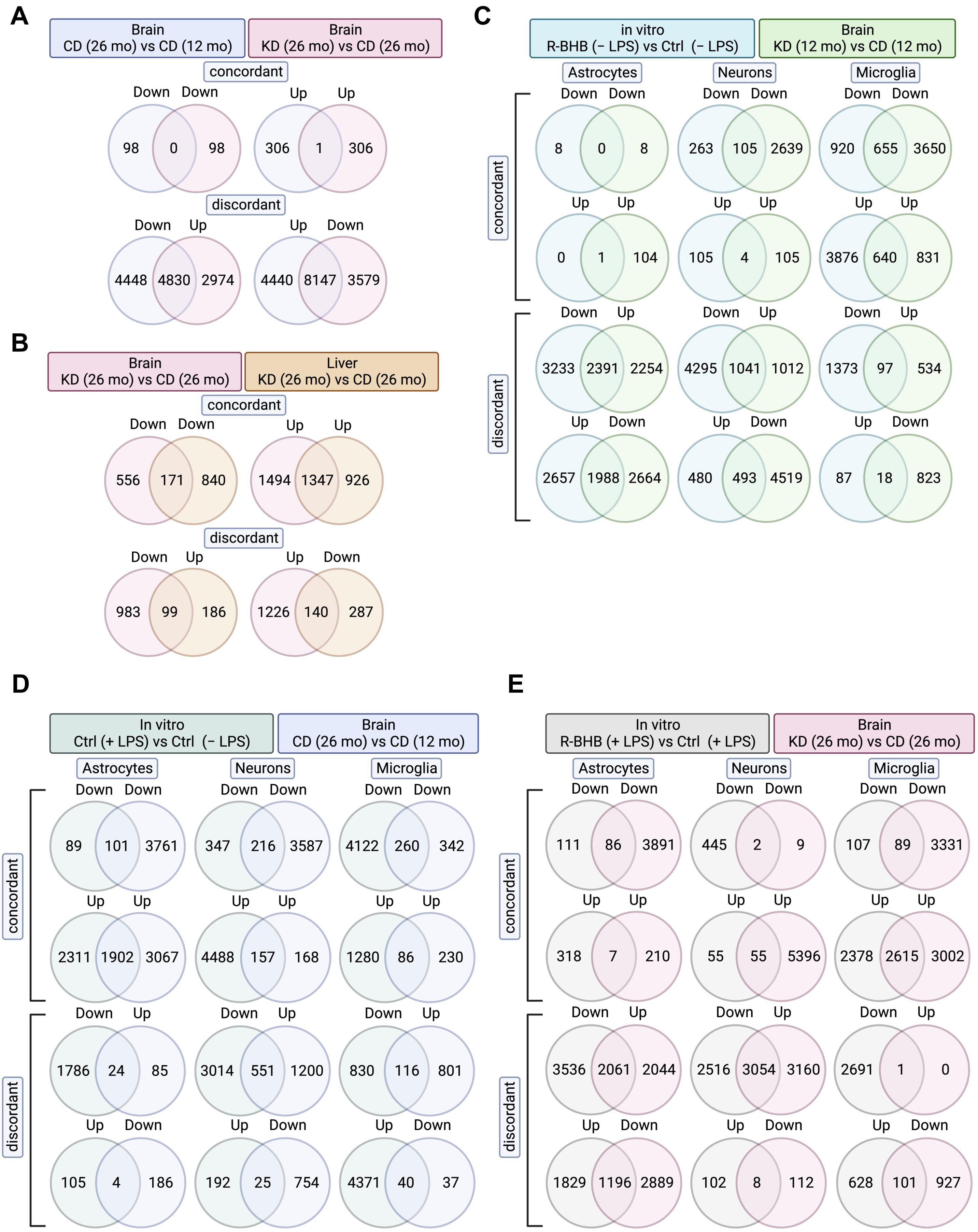
